# A single conserved residue in FlhA couples export gate activation to substrate specificity switching

**DOI:** 10.64898/2026.05.20.726703

**Authors:** Miki Kinoshita, Tohru Minamino, Motoshi Sakai, Norihiro Takekawa, Takayuki Uchihashi, Akio Kitao, Katsumi Imada, Keiichi Namba

## Abstract

FlhA is a core component of the bacterial flagellar type III secretion system that functions as both an export engine and a switch regulator of substrate specificity. How these two processes are mechanistically coupled remains unclear. Here we identify a single conserved residue in FlhA, Arg-391, as a key molecular node that links export gate activation to substrate specificity switching. The R391A substitution blocks the transition from hook-type to filament-type secretion, resulting in polyhooks lacking filaments. An intermolecular salt-bridge formation of Arg-391 with Glu-351 of an adjacent FlhA subunit stabilizes the filament-type FlhA ring. Suppressor mutations in FlhA and the ruler protein FliK restore this interaction, indicating that FliK promotes the intermolecular salt-bridge formation during export switching. Moreover, Arg-391 is required for efficient ATPase-independent activation of the export gate. Together, these findings reveal an electrostatic mechanism by which FlhA coordinates export gate activation and substrate specificity switching during flagellar assembly.

## Introduction

The bacterial flagellum is a sophisticated rotary nanomachine that enables motility under a variety of environmental conditions. Flagellar assembly proceeds in a stepwise manner, requiring the precise export of structural subunits through the flagellar type III secretion system (fT3SS). The fT3SS of *Salmonella enterica* serovar Typhimurium (hereafter referred to as *Salmonella*) transports structural subunits in a defined temporal order: rod-type proteins (FliE, FlgB, FlgC, FlgF, FlgG, and FlgJ) are secreted first, followed by hook-type proteins (FlgD, FlgE, and FliK), and finally filament-type proteins (FlgK, FlgL, FlgM, FliC, and FliD)^1^.

To efficiently build the flagellum on the cell surface, the fT3SS exhibits at least two distinct substrate specificities during assembly: one for rod-hook-type substrates and another for filament-type substrates^2,3^. During hook assembly, the fT3SS transports the FliK ruler protein, which monitors hook length and switches the substrate specificity of the fT3SS from the rod-hook-type to the filament-type once the hook reaches its mature length of approximately 55 nm in *Salmonella*^4–6^. FliK transmits the hook-length signal to the fT3SS components FlhB and FlhA, thereby terminating hook assembly and initiating filament formation^7–12^. Despite extensive studies, our understanding on the molecular mechanisms that coordinate this highly ordered assembly process remains incomplete.

The fT3SS consists of a transmembrane export gate complex comprising FlhA, FlhB, FliP, FliQ, and FliR and a cytoplasmic ATPase complex composed of FliH, FliI, and FliJ^15,16^. The export gate complex utilizes proton motive force across the cytoplasmic membrane as the energy source to drive flagellar protein export^17,18^. ATP hydrolysis by the cytoplasmic ATPase complex enables the export gate to serve as a proton-protein antiporter that couples inward-directed H^+^ flow with outward-directed protein export^19^. Furthermore, the export gate alone can autonomously function as an active protein transporter when the external Na⁺ concentration or membrane potential increases, even in the absence of the ATPase complex. This suggests that the export gate intrinsically functions as both a membrane voltage sensor and a Na⁺-driven export engine in addition to its H⁺-driven engine. The N-terminal transmembrane domain of FlhA (residues 1–327) serves as this dual-fuel export engine equipped with the voltage sensor^21–23^. The C-terminal cytoplasmic domains of FlhA (FlhA_C_) and FlhB (FlhB_C_) project into the central cavity of the C-ring, forming a substrate-docking platform for flagellar export chaperones and export substrates^23–30^. This docking platform requires the cytoplasmic ATPase complex to ensure the correct temporal order of protein export during flagellar assembly^31,32^.

FlhA_C_ (residues 328–692) consists of four domains—D1 (residues 362–434 and 484–503), D2 (residues 435–483), D3 (residues 504–583), and D4 (residues 584–692)—and a flexible linker (FlhA_L_, residues 328–361) (PDB ID: 3A5I) (Supplementary Fig. 1a)^33^. It forms a nonameric ring through intermolecular interactions between the D1-D1 and D1-D3 domains^34,35^. The D1 domain contains a positively charged cluster formed by Arg-388, Arg-391, Lys-392, and Lys-393, which interacts with Glu-351 and Asp-356 in the C-terminal region of FlhA_L_ (FlhA_L-C_) of the adjacent FlhA_C_ subunit (Fig. 1a)^33,35^. The R391A/K392A/K393A triple mutation (hereafter referred to as AAA) disrupts ring formation of FlhA_C_ in solution and inhibits switching of substrate specificity from rod-hook-type to filament-type, resulting in the formation of polyhooks without filaments^36–38^. These findings suggest that the positively charged cluster is directly involved in the FliK-induced structural transition of the FlhA_C_ ring from the rod-hook-type to the filament-type state. Furthermore, a conserved GYXLI motif (residues 368–372) in domain D1 of FlhA_C_ is required not only for cyclic domain motions during flagellar protein export^39^ but also for structural transition of the FlhA_C_ ring responsible for the substrate specificity switch^32^. However, the molecular determinants within FlhA_C_ that couple flagellar protein export activity with substrate specificity switching have remained elusive.

**Fig. 1.**
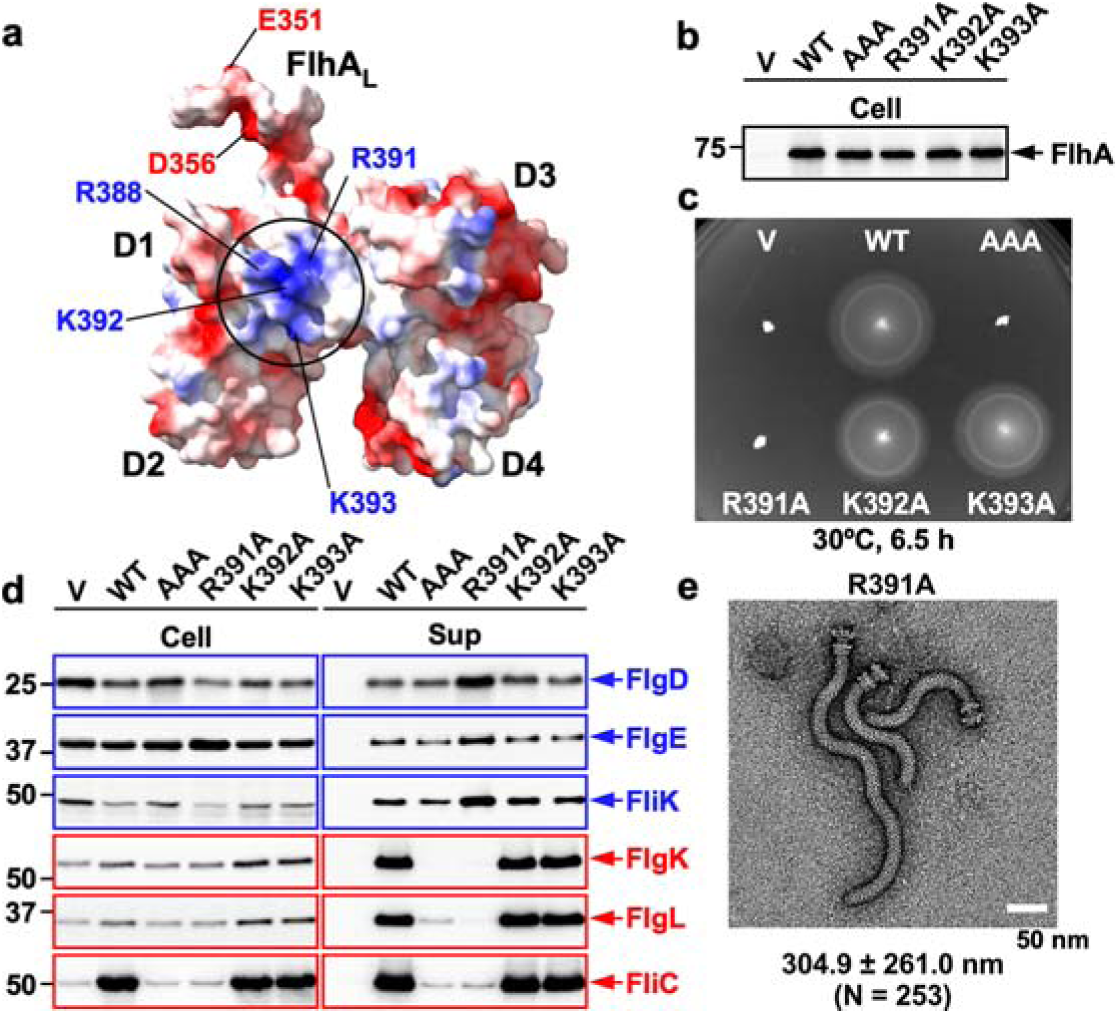
Mutational analysis of a positively charged cluster in FlhA. **(a)** Electrostatic surface potential of the C-terminal cytoplasmic domain of FlhA (FlhA_C_) (PDB ID: 3A5I). FlhA_C_ consists of four domains (D1–D4) and a flexible linker (FlhA_L_). Conserved Arg-388, Arg-391, Lys-392, and Lys-393 form a positively charged cluster on the surface of domain D1. In the 3A5I crystal structure, Glu-351 and Asp-356 in FlhA_L_ interact with the positively charged cluster of an adjacent FlhA_C_ subunit. **(b)** Effect of alanine substitutions on the steady-state expression level of FlhA. Whole-cell proteins (Cell) were prepared from the *Salmonella* NH001 (Δ*flhA*) strain carrying pTrc99AFF4 (V), pMM130 (WT), pYI004 (AAA), pMKM130(R391A), pMKM130(K392A), or pMKM130(K393A). Samples normalized to an optical density of OD_600_ were analyzed by SDS–PAGE and immunoblotting using a polyclonal anti-FlhA_C_ antibody. **(c)** Soft-agar motility assays of the transformants shown in (b). Plates were incubated at 30 °C for 6.5 h. At least seven independent assays were performed. **(d)** Secretion assays of FlgD, FlgE, FliK, FlgK, FlgL, and FliC. Whole-cell proteins (Cell) and culture supernatants (Sup) were prepared from the transformants shown in (b) and analyzed by immunoblotting using the indicated polyclonal antibodies. Hook-type and filament-type substrates are highlighted in blue and red, respectively. Molecular mass markers (kDa) are shown on the left. At least three independent assays were performed. **(e)** Electron micrograph of polyhook–basal bodies isolated from the *flhA*(R391A) mutant. The average polyhook length and standard deviation are shown. *N* indicates the number of structures measured.

Arg-391, Lys-392, and Lys-393 are highly conserved across bacterial species (Supplementary Fig. 1b), indicating strong evolutionary conservation and functional importance. To investigate the functional roles of these positively charged residues in flagellar protein export and substrate specificity switching, we analyzed the structural remodeling mechanism of the FlhA_C_ ring by mutational analysis, GST affinity chromatography, high-speed atomic force microscopy (HS-AFM), X-ray crystallography, and molecular dynamics (MD) simulations. We show that Arg-391 of FlhA plays dual and essential roles in the fT3SS. The R391A substitution alone abolishes the substrate specificity switch from rod-hook-type to filament-type secretion, resulting in the formation of polyhooks lacking filaments. Structural and biophysical analyses demonstrate that Arg-391 forms an intermolecular salt bridge with Glu-351 of the adjacent FlhA subunit, stabilizing the filament-type conformation of the FlhA_C_ ring. Suppressor mutations in FlhA and FliK restore switching function, indicating that FliK promotes this intermolecular salt-bridge formation during the transition to filament-type export. Moreover, under ATPase-deficient conditions, the R391A mutation impairs hook-type protein export, suggesting that Arg-391 also facilitates FlhA-mediated protein secretion in cooperation with FliH and FliI. These findings reveal that this single conserved residue of FlhA coordinates substrate specificity switching and protein export, providing mechanistic insight into how the fT3SS achieves temporal control during flagellar assembly.

## Results

### A positively charged cluster in FlhA_C_ is essential for substrate specificity switching

To define the functional role of the positively charged cluster comprising Arg-391, Lys-392, and Lys-393 in FlhA_C_, we individually replaced each residue with alanine and assessed motility on soft agar. Immunoblotting confirmed that these substitutions did not alter the steady-state level of FlhA (Fig. 1b). The R391A mutant was non-motile, similar to the AAA mutant, whereas the K392A and K393A mutants remained motile (Fig. 1c), indicating that Arg-391 is specifically required for FlhA function.

We next examined secretion of hook-type substrates (FlgD, FlgE, and FliK) and filament-type substrates (FlgK, FlgL, and FliC) by immunoblotting. As reported previously^36^, the AAA mutant secreted hook-type substrates at near wild-type levels but failed to secrete filament-type substrates (Fig. 1d). Notably, the R391A mutant secreted substantially higher amounts of FlgD, FlgE, and FliK than the AAA mutant, producing markedly longer polyhooks (304.4 ± 261.0 nm, N = 253) than those formed by the AAA mutant (192.5 ± 94.6 nm, N = 309) (Fig. 1e and Supplementary Fig. 2). In contrast, the R391A substitution completely abolished filament-type secretion (Fig. 1d), resulting in the absence of filaments as observed by fluorescence microscopy (Supplementary Fig. 3). Consistent with this defect, GST pull-down assays showed that the R391A substitution substantially reduced binding of FlhA_C_ to the FlgN-FlgK chaperone-filament-type substrate complex (Fig. 2a). Neither the K392A nor the K393A mutation affected secretion of either substrate class (Fig. 1d). Together, these results indicate that Arg-391 within the positively charged cluster of FlhA_C_ is the one that is essential for substrate specificity switching of the fT3SS.

**Fig. 2.**
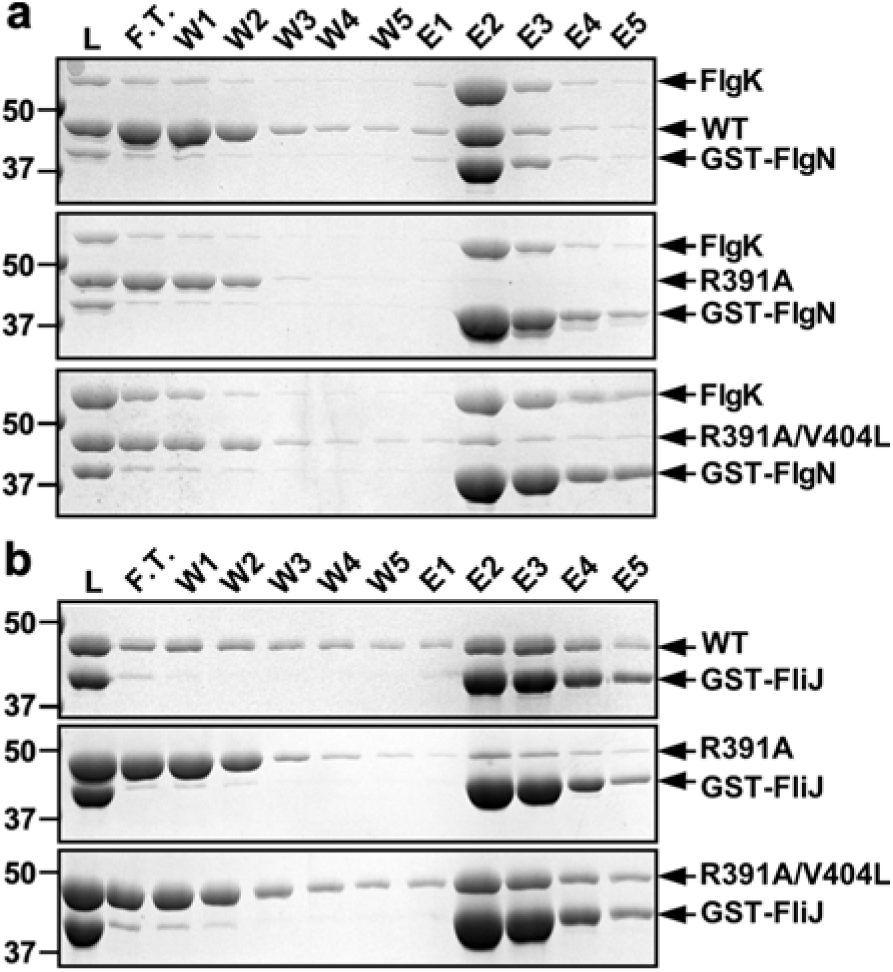
Effect of R391A and R391A/V404L mutations on interactions of FlhA_C_ with (a) the FlgN–FlgK chaperone–substrate complex and (b) FliJ. Purified His-tagged FlhA_C_ (WT, first row), His- FlhA_C_(R391A) (second row), or His-FlhA_C_(R391A/V404L) (third row) was mixed with GST-FlgN in complex with FlgK (a) or GST-FliJ (b). After overnight dialysis against PBS, samples were subjected to GST affinity chromatography. Flow-through (F.T.), wash (W), and elution (E) fractions were analyzed by Coomassie Brilliant Blue staining. Molecular mass markers (kDa) are shown on the left. Three independent assays were performed.

To understand why the AAA mutant forms shorter polyhooks than the R391A mutant, we introduced the Δ*fliK*::*tetRA* allele into the AAA background. The resulting strain produced polyhooks (205.6 ± 99.5 nm, N = 126) comparable in length to those of the AAA mutant (Supplementary Fig. 2). Because the Δ*fliK*::tetRA strain itself forms much longer polyhooks (423.6 ± 280.8 nm, N = 332), the AAA mutation likely reduces the secretion rate of hook-type substrates, thereby slowing polyhook elongation. Furthermore, replacement of Arg-391, Lys-392, and Lys-393 with acidic residues (DDD or EEE) impaired export of both FlgD and FliC, resulting in a non-motile phenotype (Supplementary Fig. 4). These observations demonstrate that the positively charged cluster in FlhA_C_ contributes not only to substrate specificity switching but also to efficient flagellar protein export.

### The positively charged cluster is required for ATPase-independent activation of the export gate

We next examined whether the positively charged cluster of FlhA contributes to flagellar protein export independently of the cytoplasmic ATPase complex. For this purpose, we used a *Salmonella* Δ*fliH-fliI flhB*(P28T) Δ*flhA* strain as the host, because the *flhB*(P28T) mutation markedly increases the frequency of flagellar assembly in the absence of FliH and FliI^17^. Flagella-driven motility and fT3SS-mediated protein secretion were assessed by soft-agar motility assays and immunoblotting using a polyclonal anti-FlgD antibody, respectively.

The K392A and K393A substitutions, which did not affect flagella-driven motility in the presence of FliH and FliI (Fig. 1c), considerably reduced the motility under ATPase-deficient conditions (Fig. 3a), indicating that FlhA variants carrying these substitutions require FliH and FliI to fully exert their protein export activity. The R391A substitution severely impaired secretion of the hook-type substrate FlgD in the absence of FliH and FliI (Fig. 3b). Similar secretion defects were observed in the AAA, K392A, and K393A mutants in their absence. In contrast, the *flhB*(P28T) mutation alone did not affect either motility or flagellar protein export (Fig. 3c,d).

**Fig. 3.**
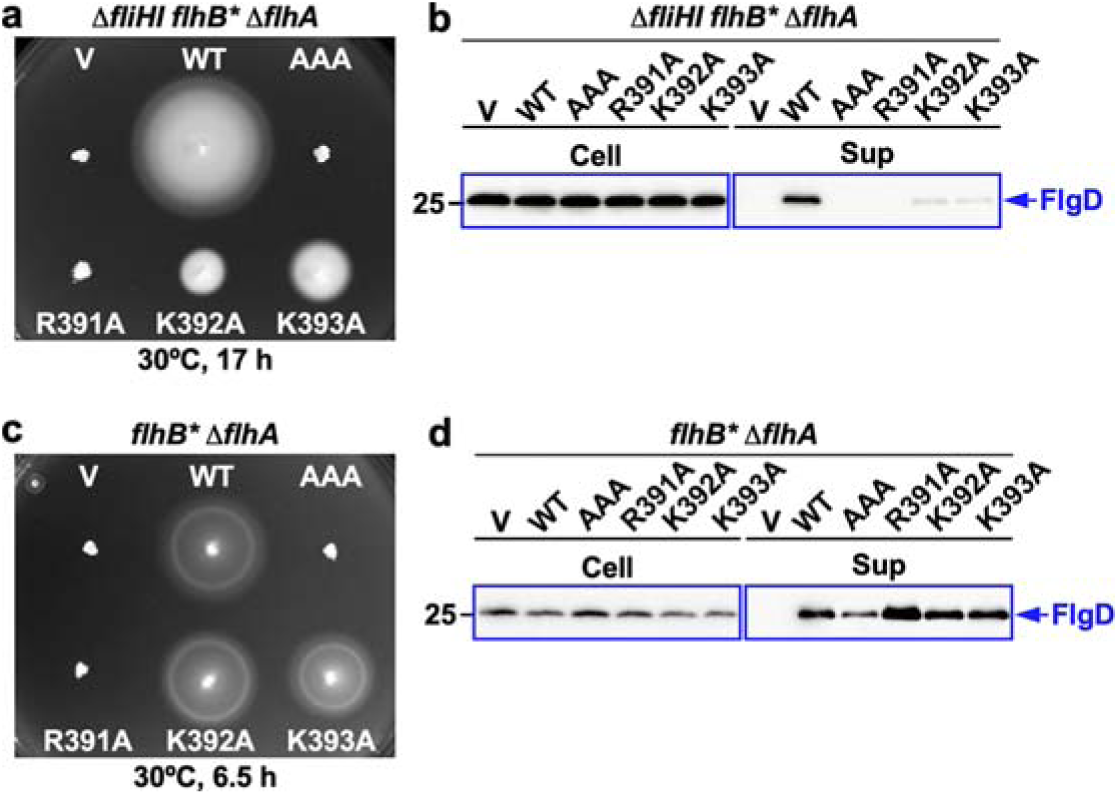
Effect of FliH and FliI deletion on motility and flagellar protein export in *flhA* alanine mutants. (a,c) Soft-agar motility assays of (a) the *Salmonella* NH004 (Δ*fliHI flhB*\* Δ*flhA*) or (c) NH002 (*flhB*\* Δ*flhA*) strain carrying pTrc99AFF4 (V), pMM130 (WT), pYI004 (AAA), pMKM130(R391A), pMKM130(K392A), or pMKM130(K393A). Plates were incubated at 30°C. At least seven independent assays were performed. **(b,d)** Immunoblot analysis of hook-type substrate secretion using a polyclonal anti-FlgD antibody. Whole-cell proteins (Cell) and culture supernatants (Sup) were prepared from the transformants shown in (a) or (c). Molecular mass markers (kDa) are indicated on the left. At least three independent assays were performed.

Notably, the secretion defects caused by substitutions in the positively charged cluster were more pronounced in the ATPase-deficient background than under ATPase-proficient conditions, indicating that this cluster is collectively required for ATPase-independent activation of the export gate. Together, these results demonstrate that the conserved positively charged cluster centered on Arg-391 contributes not only to substrate specificity switching but also to efficient activation of the flagellar export gate when the assistance by the cytoplasmic ATPase complex is unavailable.

### Isolation and characterization of suppressor mutations that bypass the R391A defect

To investigate whether the defects in FlhA caused by the R391A substitution can be bypassed under ATPase-proficient conditions, we isolated 22 up-motile suppressor mutants from the *flhA*(R391A) strain (Fig. 4a and Supplementary Fig. 5). All suppressor mutants restored secretion of filament-type substrates (FlgK and FliC) to a considerable degree, while secretion of rod/hook-type substrates (FlgD, FlgE, and FliK) was reduced compared with the parental R391A mutant (Fig. 4b). As a result, these suppressor mutants produced filaments (Supplementary Fig. 3), and their hooks were substantially shorter than those of the R391A mutant and only slightly longer than wild-type hooks (Fig. 4c).

**Fig. 4.**
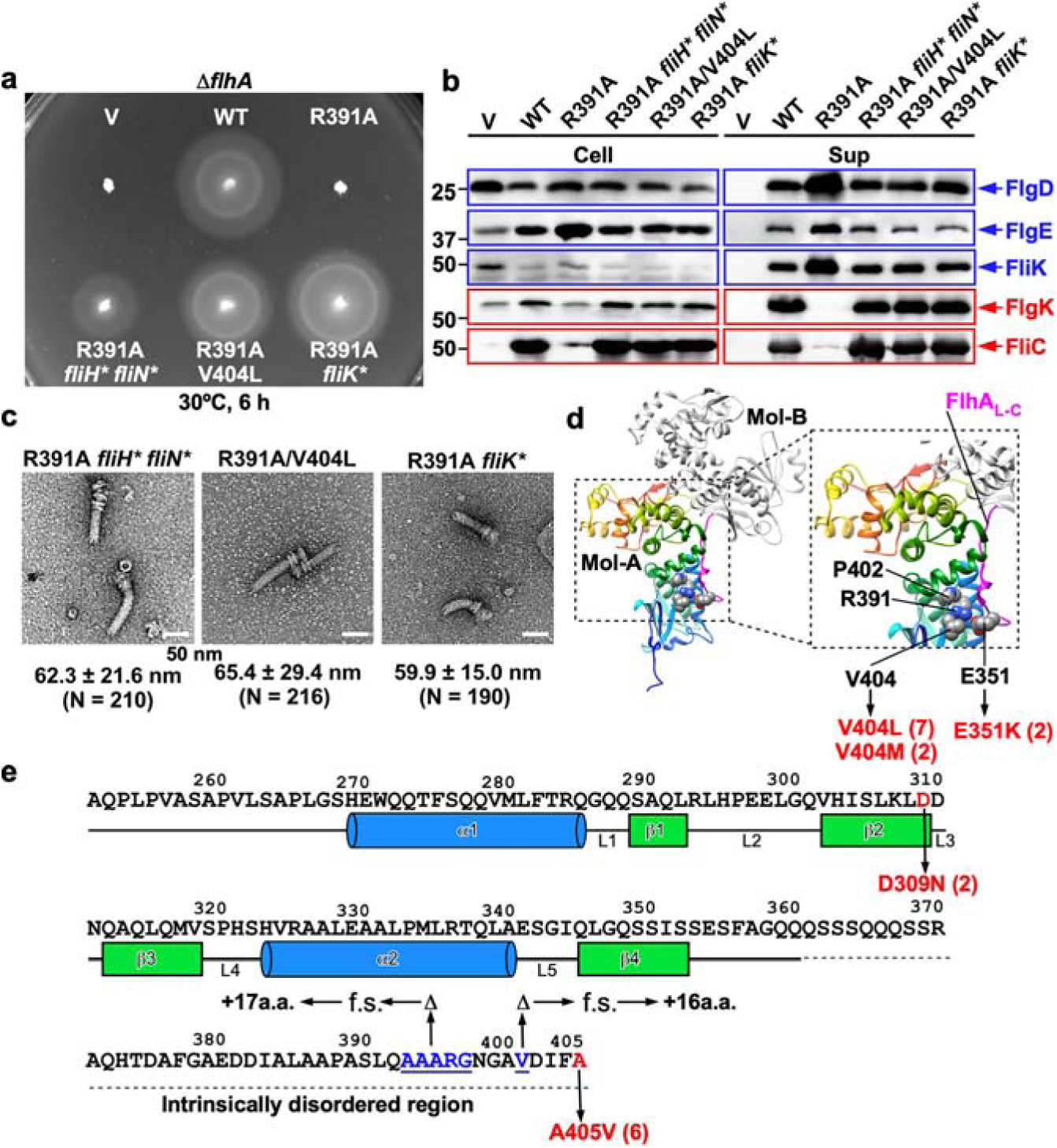
Isolation and characterization of suppressor mutations of the *flhA*(R391A) mutant. **(a)** Soft-agar motility assays of the *Salmonella* NH001 (Δ*flhA*) strain carrying pTrc99AFF4 (V), pMM130 (WT), or pMKM130(R391A) and *flhA*(R391A) suppressor strains. MMA391-6 (R391A *fliH*\* *fliN*\*) and MMA391-8 (R391A *fliK*\*) carry chromosomal suppressor mutations, whereas MMA391-7 carries an intragenic suppressor mutation [*flhA*(R391A/V404L)] on the plasmid. Plates were incubated at 30°C for 6 hours. At least seven independent measurements were performed. **(b)** Secretion assays of hook-type and filament-type substrates. Whole-cell proteins (Cell) and culture supernatants (Sup) were prepared from the strains shown in (a) and analyzed by immunoblotting using the indicated polyclonal antibodies. Hook-type and filament-type substrates are highlighted in blue and red, respectively. Molecular mass markers (kDa) are shown on the left. Three independent assays were performed. **(c)** Electron micrographs of hook–basal bodies isolated from the R391A *fliH*\* *fliN*\*, R391A/V404L, and R391A *fliK*\* strains. Average hook lengths and standard deviations are shown. *N* indicates the number of structures measured. **(d)** Location of intragenic suppressor mutations in the 3A5I structure of FlhA_C_. Two FlhA_C_ molecules, Mol-A (rainbow color) and Mol-B (light grey), are contained in the asymmetric unit of the crystal belonging to a tetragonal space group *I*4_1_. Arg-391 of Mol-A forms an intermolecular salt bridge with Glu-351 in the C-terminal linker region of FlhA (FlhA_L-C_, magenta) in Mol-B. The hydrophobic moiety of Arg-391 also makes intramolecular hydrophobic contacts with Pro-402 and Val-404, thereby stabilizing the salt bridge formation. Intragenic suppressor mutations E351K, V404L, and V404M are highlighted in red. The numbers in parentheses indicate the number of isolated mutations. **(e)** Location of extragenic suppressor mutations in the C-terminal domain of FliK (FliK_C_). FliK_C_ consists of a compactly folded core domain (residues 268–352) and an intrinsically disordered C-terminal region (residues 353–405). Secondary structure elements are shown below the amino acid sequence. The extragenic suppressor mutations D309N and A405V are highlighted in red, whereas deletions highlighted in blue cause frameshifts (f.s.), results in the addition of extra amino acids (+a.a.). The numbers in parentheses indicate the number of isolated mutations.

P22-mediated transduction and DNA sequencing revealed that suppressor mutations are located either within plasmid-borne *flhA* or in other flagellar genes on the chromosome (Fig. 4d,e). First, three missense mutations in FlhA_C_—E351K (two isolates), V404L (seven isolates), and V404M (two isolates)—were repeatedly recovered. Second, a double mutation, *fliH*(V25A) *fliN*(N24S), was identified. Third, multiple mutations were found in the C-terminal export-switch domain of FliK (FliK_C_), including two missense substitutions, D309N (two isolates) and A405V (six isolates), and two C-terminal frameshift mutations that result in the addition of extra amino acids at the C terminus.

The crystal structure of wild-type FlhA_C_ (PDB ID: 3A5I) shows that Arg-391 forms an intermolecular salt bridge with Glu-351 in FlhA_L-C_ of the adjacent FlhA_C_ subunit, stabilized by hydrophobic contacts involving Val-404 (Fig. 4d). Accordingly, the E351K, V404L, and V404M substitutions likely compensate for the loss of the positive charge and hydrophobic moiety of Arg-391 to restore the intermolecular interactions of domain D1 with FlhA_L-C_ of the adjacent FlhA_C_ subunit. GST pull-down assays demonstrated that the V404L substitution substantially restored binding of FlhA_C_(R391A) to the FlgN-FlgK chaperone-filament-type substrate complex (Fig. 2a), indicating that proper intermolecular interactions within the FlhA_C_ ring are required for formation of the filament-type chaperone-binding site.

The cytoplasmic ATPase ring complex is composed of 12 FliH subunits, 6 FliI subunits, and a single FliJ subunit. FliJ penetrates the central pore of the FliI hexamer. The FliH dimer binds to each FliI subunit through the C-terminal domain of FliH and the N-terminal domain of FliI. The FliI_6_-FliJ complex firmly anchors to the basal body C-ring through interactions between the extreme N-terminal region of FliH and a C-ring protein, FliN^15^. GST pull-down assays further showed that the R391A substitution weakens the interaction between FlhA_C_ and FliJ, whereas the V404L suppressor markedly restores this interaction (Fig. 2b). Given that the FliH_12_-FliI_6_-FliJ_1_ complex promotes structural remodeling of the FlhA_C_ ring required for substrate specificity switching^32.36^, the *fliH*(V25A) *fliN*(N24S) suppressor, located at the connecting interface between the ATPase ring complex and C-ring^40–43^, likely enhances productive engagement of FliJ with FlhA.

FliK_C_ consists of a folded core domain (residues 268–352) followed by an intrinsically disordered region (residues 353–405), with residues 301–350 and the terminal five residues (401–405) being essential for export switching^10^. AlphaFold3-based structural predictions suggest direct contacts between FliK_C_ and FlhA_C_ (Supplementary Fig. 6). In this model, the terminal five residues of FliK form an antiparallel five-stranded β-sheet with four β-strands in the D1 domain of FlhA_C_, and Ala-405 of FliK makes a hydrophobic contact with helix α2 in domain D1 of FlhA_C_, which harbors the conserved positively charged cluster. The conserved GYXLI motif, previously shown to be required for FlhA_C_ ring remodeling^32^, is predicted to interact with the FliK_C_ core. Consistent with this model, the FlhA(Y369A/R370A/L371A/I372A) mutant exhibits a polyhook-filament phenotype, whereas the A405V substitution in FliK_C_ also restores the export switching function of this FlhA mutant to a considerable degree. This suppression not only shortens the polyhook length to nearly that of the wild type but also enables filament formation^32^. Together, these findings support a model in which the interaction of FliK_C_ with FlhA_C_ induces conformational remodeling of the D1 domain of FlhA_C_, enabling Glu-351 and Asp-356 in FlhA_L-C_ to engage the positively charged cluster in the adjacent FlhA subunit and trigger dissociation of FliK_C_ upon hook completion.

### Side-chain interactions around the Arg-391-Glu-351 interface coordinate ATPase-independent activation and substrate switching

The *flhA*(V404M) mutation was originally isolated as a gain-of-function allele in a *Salmonella* Δ*fliH* and Δ*fliH-fliI* backgrounds^17,44^. We therefore first examined whether substitutions around the Arg-391-Glu-351 interface activate the export gate in the absence of the cytoplasmic ATPase complex. For this purpose, we introduced a plasmid encoding the *flhA*(E351K) or *flhA*(V404L) allele into Δ*fliH-fliI* Δ*flhA* and Δ*fliH-fliI flhB(P28T)* Δ*flhA* strains and performed soft-agar motility assays and flagellar secretion assays.

The V404L substitution alone bypassed the requirement for FliH and FliI, enhancing motility (Fig. 5a,c) and secretion of both the hook-type substrate FlgD and the filament-type substrate FliC (Fig. 5b,d), similarly to the previously characterized V404M mutation. In contrast, the E351K substitution inhibited motility in the ATPase deficient background (Fig. 5a,c). The E351K substitution increased FlgD secretion compared to the wild-type allele but reduced FliC secretion (Fig. 5b,d). This suggests that electrostatic repulsion between Arg-391 and Lys-351 strongly affects efficient transition to the filament-type state of the FlhA_C_ ring.

**Fig. 5.**
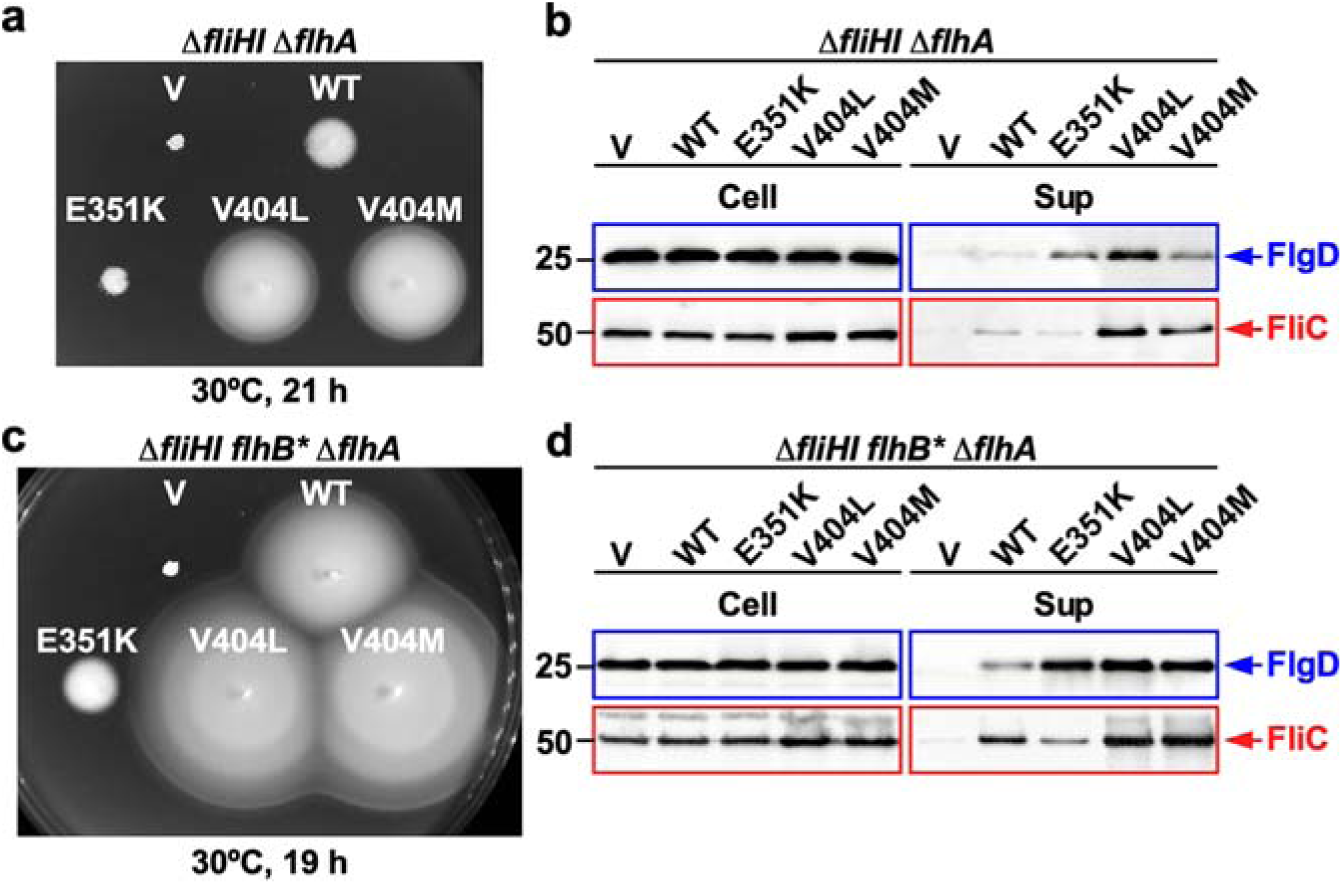
Bypass effects of intragenic suppressor mutations on FliH–FliI deficiency. (a,c) Soft-agar motility assays of the *Salmonella* (a) NH003 (Δ*fliHI* Δ*flhA*) or (c) NH004 (Δ*fliHI flhB*\* Δ*flhA*) strain carrying pTrc99AFF4 (V), pMM130 (WT), pMM130(E351K) (indicated as E351K), pMKM130(V404L), or pMKM130(V404M). Plates were incubated at 30°C. At least seven independent assays were performed. **(b,d)** Immunoblot analysis of hook-type (FlgD) and filament-type (FliC) substrate secretion. Whole-cell proteins (Cell) and culture supernatants (Sup) were prepared from the transformants shown in (a) or (c). Hook-type and filament-type substrates are highlighted in blue and red, respectively. Molecular mass markers (kDa) are shown on the left. Three independent assays were performed.

Notably, although these suppressor mutations enhanced hook-type secretion in the ATPase-deficient background, combining either mutation with *flhA*(R391A) abolished both flagella-driven motility and flagellar protein export in the absence of FliH and FliI (Supplementary Fig. 7). These results indicate that productive activation of FlhA requires not merely the presence of charged residues at this interface but their correct electrostatic configuration and identify Arg-391 as a central determinant that integrates ATPase-independent export gate activation with subsequent substrate specificity switching.

### Intermolecular salt-bridge formation stabilizes the filament-type FlhA_C_ ring

HS-AFM and mutational analyses have previously shown that the interactions of FlhA_L-C_ with the D1 and D3 domains of an adjacent FlhA_C_ subunit stabilize the filament-type FlhA_C_ ring structure on mica surface^32,36^. To obtain direct evidence that intermolecular salt-bridge formation is required for efficient and robust formation of the filament-type ring with a properly formed chaperone-binding site, we purified N-terminally His-tagged FlhA_C_(R391A) and FlhA_C_(R391A/V404L) monomers by size-exclusion chromatography (Supplementary Fig. 8a) and analyzed their ring-forming ability by HS-AFM.

HS-AFM imaging revealed that the FlhA_C_(R391A) monomer failed to form stable ring-like assemblies on the mica surface. Instead, individual particles remained largely dispersed, consistent with the inability of the R391A mutant to engage Glu-351 in FlhA_L-C_ of the adjacent subunit. In contrast, FlhA_C_(R391A/V404L) exhibited transient ring formation in HS-AFM. Although ring-like structures were detectable, they were highly unstable and rapidly disassembled. Because far-UV circular dichroism measurements indicated that the R391A and R391A/V404L substitutions did not severely perturb the overall FlhA_C_ fold (Supplementary Fig. 8b), these results suggest that the V404L substitution only partially compensates for the loss of the E351-R391 interaction caused by the R391A mutation (Fig. 6a).

**Fig. 6.**
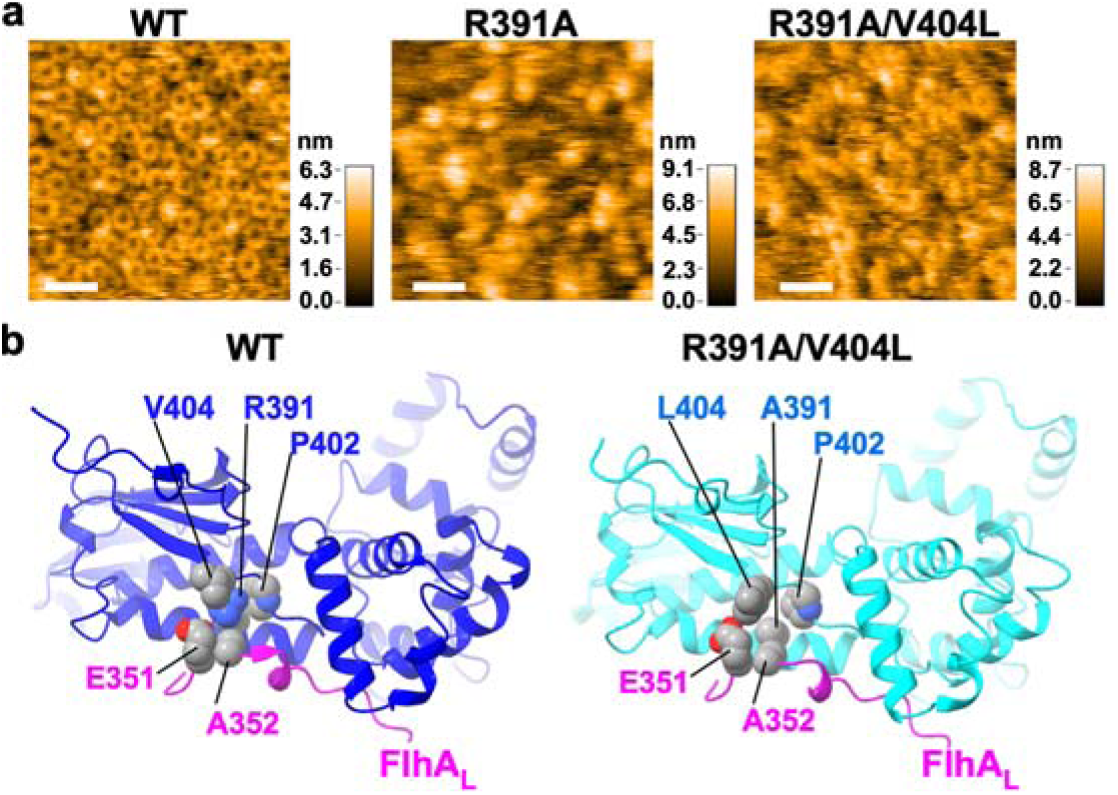
Effect of R391A and R391A/V404L mutations on intermolecular salt bridge formation between Arg-391 and Glu-351. **(a)** Representative HS-AFM images of His- FlhA_C_ (WT), His- FlhA_C_(R391A), and His- FlhA_C_(R391A/V404L) on a mica surface at a protein concentration of 2 μM. Images were recorded at 500 ms per frame over a scanning area of 150 × 150 nm² with a resolution of 150 × 150 pixels. Color bars indicate particle height (nm). Scale bars represent 30 nm. **(b)** Crystal structures of the open forms of wild-type FlhA_C_ (PDB ID: 3A5I, blue) and FlhA_C_(R391A/V404L) (PDB ID: 23PK, cyan). In the wild-type structure, Arg-391 forms an intermolecular salt bridge with Glu-351 in the linker region of an adjacent FlhA subunit (FlhA_L_, magenta). The hydrophobic moiety of Arg-391 also mediates intra-and intermolecular hydrophobic contacts with Ala-352, Pro-402, and Val-404. In the R391A/V404L structure, Leu-404 partially compensates for the loss of Arg-391 by forming hydrophobic contacts that allow FlhA_L_ to associate with the adjacent subunit, although these interactions are weaker than in the wild-type structure.

The wild-type *Salmonella* FlhA_C_ structure adopts an open conformation in a tetragonal *I*4_1_ crystal (PDB ID: 3A5I)^33^ and a semi-closed conformation in an orthorhombic *P*2_1_2_1_2_1_ crystal (PDB ID: 6AI0)^37^. In the open-form structure, two molecules (Mol-A and Mol-B) are present in an asymmetric unit, and their intermolecular D1-D3, D3-D3, and FlhA_L-C_-D1 interactions are conserved in the FlhA_C_ ring structure of *Vibrio parahaemolyticus* (PDB ID: 7AMY)^35^ (Supplementary Fig. 1a). We therefore investigated how the V404L substitution partially compensates for the loss of the E351-R391 intermolecular interaction by X-ray crystallographic analysis.

Crystallization trials of FlhA_C_(R391A/V404L) yielded crystal forms corresponding to both the open and semi-closed conformations of FlhA_C_ (Supplementary Fig. 9a,b), albeit at lower resolutions (open form, 3.95 Å; semi-closed form, 2.79 Å) than those of wild-type FlhA_C_ (open form, 2.8 Å; semi-closed form, 2.4 Å)^33,37^ (Supplementary Table 1), suggesting reduced conformational stability relative to the wild type. In contrast, crystallization of FlhA_C_(R391A) did not yield crystals corresponding to the open conformation. However, an orthorhombic *P*2_1_2_1_2_1_ crystal form corresponding to the semi-closed conformation was obtained (Supplementary Fig. 9b), although the resolution (3.2 Å) was substantially lower than that of wild-type FlhA_C_ (Supplementary Table 1). Together, these results indicate that Arg-391 contributes to stabilizing both the open and semi-closed conformations of FlhA_C_.

Structural comparison further revealed that, in the R391A/V404L mutant, FlhA_L-C_ still associates with the adjacent FlhA_C_ subunit, but the intermolecular hydrophobic contacts are weakened relative to the wild type (Fig. 6b). This reduced interaction strength provides a structural explanation for the instability of the filament-type ring observed by HS-AFM. Collectively, these results demonstrate that the E351-R391 intermolecular salt bridge is a key stabilizing element of the filament-type FlhA_C_ ring and that partial restoration of hydrophobic interactions can reconstitute the filament-type conformation, albeit in a less stable form.

To examine whether these structural differences are dynamically maintained in solution, we performed all-atom MD simulations of FlhA_C_(R391A) and FlhA_C_(R391A/V404L) at 27°C for 8 μs. Because the transition from the open to the closed conformation involves hinge motions between domains D1-D3 and D3-D4, bringing domain D4 closer to domain D2^33,37^, we calculated the center-of-mass distance (*d*_24_) between domains D2 and D4 throughout the simulations. When simulations were initiated from the open conformations, both FlhA_C_(R391A) and FlhA_C_(R391A/V404L) initially exhibited cyclic open-close domain motions, similar to those observed for wild-type FlhA_C_^33,37^ (Supplementary Fig. 10a). However, unlike FlhA_C_(R391A/V404L), FlhA_C_(R391A) ceased this dynamic behavior after approximately 3.6 μs and adopted a closed conformation.

We next compared representative MD-derived structures with the atomic structure of FlhA_C_ in complex with the FliS-FliC fusion protein (PDB ID: 6CH3)^28^ (Supplementary Fig. 10b). Flagellar chaperone-substrate complexes, including FlgN-FlgK, FliS-FliC, and FliT-FliD, bind to a conserved hydrophobic dimple at the interface between domains D1 and D2 of FlhA_C_. This interaction involves highly conserved residues, including Phe-459 of FlhA_C_ and Tyr residues of the chaperones (Tyr-122 of FlgN, Tyr-10 of FliS, and Tyr-106 of FliT)^25,27^. In the R391A/V404L mutant, Tyr-10 of FliS formed a hydrophobic contact with Phe-459 in a manner similar to that observed in wild-type FlhA_C_. In contrast, this interaction was not observed in FlhA_C_(R391A). These findings are consistent with our pull-down experiments showing that the R391A mutation disrupts interaction with the FlgN-FlgK complex, whereas the V404L substitution restores this interaction to a significant extent (Fig. 2a). Together, these results suggest that the E351-R391 intermolecular salt bridge facilitates proper formation of the chaperone-docking site at the D1-D2 interface of FlhA_C_.

## Discussion

Flagellar assembly requires precise temporal coordination between flagellar protein assembly at the distal growing end and export substrate specificity switching by the fT3SS at the base. In this study, we identify a single conserved residue of FlhA, Arg-391, as a key molecular node that couples these two fundamental processes. Through a combination of mutational analysis, suppressor genetics, high-speed atomic force microscopy, X-ray crystallography, and MD simulations, we show that Arg-391 is essential not only for the transition from rod-hook-type to filament-type secretion (Fig. 1) but also for efficient activation of the export gate, particularly under conditions where the cytoplasmic ATPase complex is absent (Fig. 3).

Our data support a model in which Arg-391 mediates an intermolecular salt bridge with Glu-351 of an adjacent FlhA subunit in the FlhA_C_ ring, stabilizing the filament-type conformation. Disruption of this interaction by the R391A substitution prevents stable ring formation, as evidenced by the absence of ring structures in the HS-AFM images and the inability to obtain a 3A5I-like crystal structure. Suppressor mutations such as V404L partially restore the intermolecular interaction between FlhA_L-C_ and domain D1, allowing transient and unstable ring assembly as observed by HS-AFM (Fig. 6), which is also consistent with the reduced crystallographic resolution of the FlhA_C_ structure. Furthermore, MD simulations revealed that the R391A mutation inhibits proper formation of the chaperone-docking site at the interface between domains D1 and D2 of FlhA_C_, whereas V404L restores this formation (Supplementary Fig. 10). These results indicate that formation of the Arg-391-Glu-351 intermolecular salt bridge is a structural prerequisite for establishing the filament-type state of the FlhA_C_ ring. This is consistent with our observations that the R391A mutation inhibits the interactions of FlhA_C_ with the FlgN-FlgK chaperone-filament-type substrate complex and that the V404L substitution substantially restores binding of FlhA_C_(R391A) to the FlgN-FlgK complex (Fig. 2a).

Together, these observations support a model in which the filament-type FlhA_C_ ring is a dynamic assembly that requires the E351-R391 intermolecular salt bridge to maintain structural integrity for its proper function. In this model, suppressor mutations restore export activity not by reconstituting a permanently stable ring, but by shifting the equilibrium toward a sufficiently long-lived filament-type assembly that permits productive interactions with flagellar export chaperones in complex with their cognate filament-type substrates.

Importantly, our findings further reveal that the positively charged cluster centered on Arg-391 plays a critical role in ATPase-independent activation of the export gate. In the absence of FliH and FliI, substitutions within this cluster severely impaired hook-type protein export (Fig. 3), demonstrating that FlhA possesses an intrinsic activation mechanism that relies on electrostatic interactions within the entire FlhA ring. The more pronounced export defects observed under ATPase-deficient conditions suggest that the cytoplasmic ATPase complex normally compensates for weakened intermolecular interactions in the FlhA ring.

Extragenic suppressor mutations in FliK_C_, which catalyzes substrate specificity switching of the fT3SS upon hook completion^10^ (Fig. 4) provide further insight into how this switching mechanism is regulated during hook assembly. The AlphaFold3-based model of the FliK_C_-FlhA_C_ complex reveals that the residues previously implicated in polyhook formation and suppressor mutations cluster at or near the predicted FlhA_C_-FliK_C_ interface (Supplementary Fig. 6). This structural clustering of parental and suppressor mutations provides independent support for a direct role of FliK_C_ in inducing the conformational remodeling of the FlhA_C_ ring that underlies the substrate specificity switching.

Once ATP is hydrolyzed by the cytoplasmic FliH_12_-FliI_6_-FliJ_1_ ATPase ring complex, direct interactions of FliJ with FlhA_L-C_ enable the export gate to become an active H^+^-driven protein transporter^19,22^. Thus, the ATPase ring complex is directly involved in the activation of the export gate. Furthermore, this ATPase complex is also required for structural remodeling of the FlhA_C_ ring responsible for substrate specificity switching^32,36^. Here, we found that the *fliH*(V25A) *fliN*(N24S) double mutation bypasses the R391A defect (Fig. 4). Because the cytoplasmic ATPase ring complex associates with the basal body C-ring through an interaction between FliH and FliN^40–43^, the *fliH*(V25A) *fliN*(N24S) mutation likely alters the ATPase-C-ring interface. Therefore, we suggest that this double mutation affects the FlhA_L-C_-FliJ interaction. Given that the R391A mutation weakens the FlhA_L-C_-FliJ interaction, whereas the V404L suppressor mutation enhances it (Fig. 2b), we propose that FliJ plays dual roles: activating the export gate and facilitating FlhA ring remodeling required for substrate specificity switching through the interaction with FlhA_L-C_.

Together, these observations lead us to propose an integrated model in which Arg-391 acts as a molecular switch that links export gate activation to substrate specificity switching through electrostatic control of FlhA_C_ ring architecture. During hook assembly, the FliH_12_-FliI_6_-FliJ_1_ complex assists the FlhA ring in maintaining efficient H^+^-driven protein export even when intermolecular interactions between FlhA_C_ subunits are weak. Upon hook completion, the FliH_12_-FliI_6_-FliJ_1_ complex also assists FliK_C_ in promoting formation of the Arg-391-Glu-351 intermolecular salt bridge, driving the transition of the FlhA_C_ ring into the filament-type ring conformation that enforces the switch in substrate specificity. By concentrating both regulatory functions into a single conserved residue, the fT3SS achieves a robust and tightly synchronized control of flagellar assembly.

Beyond flagellar assembly, our findings have broader implications for virulence-associated type III secretion system (vT3SS) in bacterial pathogenicity, also known as injectisome. The core architecture of the fT3SS is evolutionarily related to vT3SSs, which likewise rely on a precise control of secretion activity and substrate hierarchy. FlhA is homologous to the inner membrane export gate components of vT3SSs, suggesting that electrostatic coupling between export gate activation and substrate specificity switching may represent a conserved regulatory principle^34,35^. In this context, the Arg-391-Glu-351 intermolecular salt bridge provides a paradigm for how subtle electrostatic interactions within oligomeric export platforms can integrate energy utilization, conformational remodeling, and secretion order. Given that vT3SSs also transition between early and late substrates during infection, analogous mechanisms may underlie the temporal regulation of effector secretion. Thus, our study reveals a fundamental design logic of T3SSs and provides a structural framework for understanding how bacterial nanomachines coordinate energy transduction with hierarchical protein export.

## Methods

### Bacterial strains, plasmids, P22-mediated transduction, and media

*Salmonella* strains and plasmids used in this study are listed in Table 1. To identify and purify extragenic suppressor mutations, P22-mediated transduction was performed using P22HT*int*^45^. L-broth contained 10 g of Bacto-Tryptone, 5 g of yeast extract and 5 g of NaCl per liter. Soft tryptone agar plates contained 10 g of Bacto Tryptone, 5 g of NaCl and 0.35% (w/v) Bacto-Agar per liter. Ampicillin and tetracycline were added as needed at a final concentration of 100 μg ml^-1^.

### Sequence conservation analysis of FlhA_C_

Evolutionarily conserved residues of FlhA_C_ were analyzed using the ConSurf web server (https://consurf.tau.ac.il/consurf_index.php)^46^.

### DNA manipulations

DNA manipulations were performed using standard protocols. Site-directed mutagenesis was carried out using Prime STAR Max Premix as described in the manufacturer’s instructions (Takara Bio). All mutations were confirmed by DNA sequencing (Eurofins Genomics).

### Soft-agar motility assays

Fresh colonies were inoculated onto soft tryptone agar plates and incubated at 30°C. At least seven measurements were carried out.

### Secretion assays

*Salmonella* cells were grown in L-broth supplemented with ampicillin at 30°C with shaking until the cell density reached an OD_600_ of approximately 1.2–1.4. Cultures were then centrifuged to separate the cell pellets and culture supernatants. Proteins in whole-cell and culture supernatant fractions were normalized to the OD_600_ of each culture to ensure equivalent cell numbers.

Cell pellets were resuspended directly in SDS loading buffer [62.5 mM Tris–HCl (pH 6.8), 2% (w/v) sodium dodecyl sulfate (SDS), 10% (w/v) glycerol, and 0.001% (w/v) bromophenol blue] supplemented with 1 μl of 2-mercaptoethanol. Proteins in the culture supernatants were precipitated with 10% (v/v) trichloroacetic acid, resuspended in Tris-SDS loading buffer (one volume of 1 M Tris-HCl mixed with nine volumes of SDS loading buffer) containing 1 μl of 2-mercaptoethanol, and heated at 95°C for 3 min.

Protein samples were separated by SDS–polyacrylamide gel electrophoresis (SDS-PAGE) and transferred to nitrocellulose membranes (Cytiva) using a transblotting apparatus (Hoefer). Then, immunoblotting with polyclonal anti-FlgD, anti-FlgE, anti-FliK, anti-FlgK, anti-FlgL, anti-FliC, anti-FliD, or anti-FlhA_C_ antibody as the primary antibody and anti-rabbit IgG, HRP-linked whole Ab Donkey (GE Healthcare) as the secondary antibody was carried out. Immunoblotting was performed with an iBind Flex Western Device according to the manufacturer’s instructions (Thermo Fisher Scientific). Chemiluminescent signals were detected using Amersham ECL Prime Western Blotting Detection Reagent (Cytiva) and captured with a Luminoimage Analyzer LAS-3000 (GE Healthcare). Image data were processed using Photoshop software (Adobe). The assay was performed at least three times to confirm the reproducibility of the results.

### Hook-length measurements

*Salmonella* cells were grown in 500 ml of L-broth containing ampicillin at 30°C with shaking until the cell density had reached an OD_600_ of ca. 1.0. After centrifugation (10,000 g, 10 min, 4°C), the cells were suspended in 20 ml of ice-cold sucrose buffer containing 0.1 M Tris-HCl (pH 8.0) and 0.5 M sucrose. EDTA and lysozyme were added at the final concentrations of 10 mM and 0.1 mg ml^-1^, respectively. The cell suspensions were stirred for 30 min at 4°C, and Triton X-100 and MgSO_4_ were added at final concentrations of 1% (w/v) and 10 mM, respectively. After stirring on ice for 1 hour, cell lysates were adjusted to pH 10.5 with 5 M NaOH and then centrifuged (10,000 g, 20 min, 4°C) to remove cell debris. After ultracentrifugation (45,000 g, 60 min, 4°C), pellets were resuspended in buffer A containing 10 mM Tris-HCl (pH 8.0), 5 mM EDTA, and 1% (w/v) Triton X-100, and this solution was loaded onto a 20–50% (w/w) sucrose density gradient in buffer A. After ultracentrifugation (49,100 g, 13 h, 4°C), hook-basal bodies and polyhook-basal bodies with or without filaments were collected and ultracentrifuged (60,000 g, 60 min, 4°C). Pellets were suspended in acidic buffer containing 50 mM glycine (pH 2.5) and 0.1% (w/v) Triton X100 to depolymerize flagellar filaments. After ultracentrifugation (60,000 g, 60 min, 4°C), pellets were resuspended in 50 μl of buffer A.

Samples were negatively stained with 2% (w/v) uranyl acetate. Electron micrographs were taken using JEM-1400Flash (JEOL, Tokyo, Japan) operated at 100 kV. The length of hooks and polyhooks was measured by ImageJ version 1.54p (National Institutes of Health).

### Fluorescence microscopy

*Salmonella* cells were grown in 5 ml of L-broth containing ampicillin. The cells were attached to a cover slip (Matsunami glass, Japan), and unattached cells were washed away with motility buffer containing 10 mM potassium phosphate (pH 7.0), 0.1 mM EDTA, and 10 mM sodium □-lactate. Then, flagellar filaments were labelled using anti-FliC antibody and anti-rabbit IgG conjugated with Alexa Fluor 594 (Invitrogen). After washing twice with motility buffer, the cells were observed by an inverted fluorescence microscope (IX-83, Olympus) with a 150× oil immersion objective lens (UApo150XOTIRFM, NA 1.45, Olympus) and an Electron-Multiplying Charge-Coupled Device camera (iXon^EM^+897-BI, Andor Technology). Fluorescence images of filaments labeled with Alexa Fluor 594 were merged with bright field images of cell bodies using ImageJ software version 1.54p (National Institutes of Health).

### Purification of His-tagged wild-type FlhA_C_ and its mutant variants

His-FlhA_C_ and its mutant variants were expressed in the *E. coli* BL21 Star (DE3) strain carrying a pET19b-based plasmid and purified by Ni-NTA affinity chromatography with a Ni-NTA agarose column (QIAGEN), followed by size exclusion chromatography with a HiLoad 26/600 Superdex 75 pg column (GE Healthcare) at a flow rate of 2.5 ml min^-1^ equilibrated with buffer containing 50 mM Tris-HCl (pH 8.0), 150 mM NaCl, and 1 mM EDTA.

### Analytical size exclusion chromatography

A 500 μl solution of purified His-FlhA_C_ and its mutant variants (10 μM) was applied to a Superdex 75 HR 10/300 column (GE Healthcare) equilibrated with buffer containing 50 mM Tris-HCl (pH 8.0) and 150 mM NaCl at a flow rate of 0.5 ml min^-1^. Bovine serum albumin (66.4 kDa) and ovalbumin (44 kDa) were used as size markers. All fractions were run on SDS-PAGE and then analyzed by Coomassie Brilliant blue staining.

### Far-UV circular dichroism (CD) spectroscopy

Far-UV CD spectroscopy of His-FlhA_C_ or its mutant variants was carried out at room temperature using a Jasco-720 spectropolarimeter (JASCO International Co., Tokyo, Japan). The CD spectra of His-FlhA_C_ and its mutant forms were measured in 20 mM Tris-HCl, pH 8.0 using a cylindrical fused quartz cell with a path length of 0.1 cm in a wavelength range of 200 nm to 260 nm. Spectra were obtained by averaging five successive accumulations with a wavelength step of 0.5 nm at a rate of 20 nm min^-1^, response time of 8 sec, and bandwidth of 2.0 nm.

### Pull-down assays by GST affinity chromatography

FlgK was overexpressed in *E. coli* BL21 Star (DE3) cells carrying pMKGK2 and purified by anion exchange chromatography with a Q-Sepharose high-performance column (GE Healthcare).

GST-FlgN and GST-FliJ were purified by GST affinity chromatography with a glutathione Sepharose 4B column (GE Healthcare) from the soluble fractions from the S*almonella* SJW1368 strain transformed with a pGEX-6p-1-based plasmid.

Purified His-FlhA_C_ or its mutant variants were mixed with purified GST-FlgN/FlgK complex or purified GST-FliJ, and each mixture was dialyzed overnight against PBS (8 g of NaCl, 0.2 g of KCl, 3.63 g of Na_2_HPO_4_•12H_2_O, 0.24 g of KH_2_PO_4_, pH 7.4 per liter) at 4°C with three changes of PBS. A 5 ml solution of each mixture was loaded onto a glutathione Sepharose 4B column (bed volume, 1 ml) pre-equilibrated with 20 ml of PBS. After washing of the column with 10 ml PBS at a flow rate of ca. 0.5 ml min^-1^, bound proteins were eluted with elution buffer containing 50 mM Tris-HCl (pH 8.0) and 10 mM reduced glutathione. All fractions were run on SDS-PAGE and then analyzed by Coomassie Brilliant blue staining.

### High-speed atomic force microscopy

HS-AFM imaging of FlhA_C_ ring formation was carried out in solution using a laboratory-built HS-AFM system^47,48^. A 2 μl aliquot of purified FlhA_C_ monomer (2 μM) in buffer containing 50 mM Tris-HCl (pH 8.0) and 100 mM NaCl was deposited onto a freshly cleaved mica surface mounted on a cylindrical glass stage. After incubation at room temperature for 3 min, the mica surface was rinsed thoroughly with 20 μl of 10 mM Tris-HCl (pH 6.8) to remove unbound molecules. The sample was then immersed in a liquid cell containing 60 μl of 10 mM Tris-HCl (pH 6.8).

HS-AFM imaging was carried out at room temperature in tapping mode using small cantilevers (AC7, Olympus). Images were acquired at a frame rate of 500 ms per frame over a scan area of 150 × 150 nm^2^ with a pixel resolution of 150 × 150 pixels. AFM images were processed using a laboratory-developed Python-based viewer (pyNuD) to apply plane and line-by-line slope corrections.

### X-ray crystallographic study of FlhA_C_(R391A) and FlhA_C_(R391A/V404L)

*E. coli* BL21(DE3) Star cells carrying pMKM008(R391A) or pMKM008(R391A/V404L) were inoculated into 2 l L-broth containing ampicillin and cultured at 30°C for 24 h. The cells were harvested by centrifugation (6,700 × *g* for 5 min, 4°C) and stored at -80°C. The cells were thawed, suspended in buffer A [20 mM Tris-HCl (pH 8.0), 150 mM NaCl] containing cOmplete EDTA-free (Roche) and lysozyme (Wako), and sonicated on ice. The cell lysate was centrifuged (20,400 × *g*, 15 min, 4°C) to remove cell debris. The supernatant was loaded onto a HisTrap HP 5 mL column (Cytiva) equilibrated with buffer A containing 5 mM imidazole, and bound proteins were eluted by a linear 5–500 mM gradient of imidazole in buffer A. The eluted protein solution was mixed with thrombin to cleave the N-terminal His_6_-tag and dialyzed against buffer A at 4°C for 20 h. The dialyzed protein solution was loaded onto a HisTrap HP 5 mL column (Cytiva) equilibrated with buffer A containing 20 mM imidazole, and the flow-through was collected to remove uncleaved proteins. The protein was concentrated by ultrafiltration using an Amicon Ultra-15 device (Merck Millipore). The protein sample was further purified with size exclusion chromatography using a Superdex 75 10/300□GL column (Cytiva) in buffer A. The peak fractions were collected and concentrated to 10 mg ml^-1^ using an Amicon Ultra-15 device.

Initial crystallization screening was performed by the sitting-drop vapor-diffusion technique with commercially available screening kits Wizard Classic I, II, III and IV, Wizard Cryo I and II, PEGRx I and II (Rigaku), and Crystal Screen I and II (Hampton Research) at 277K or 293K. Each drop was prepared by mixing 0.5 µl of protein solution (10 mg ml^-1^) with 0.5 µl of reservoir solution and equilibrated to 90 µl of reservoir solution. After initial screening, crystallization conditions were optimized by varying the precipitant concentration, pH, and additives using the sitting-drop vapor-diffusion method. The crystals used for the X-ray diffraction data collection were grown by using reservoir solution containing 100 mM HEPES-NaOH (pH7.5), 50 mM Li_2_SO_4_, 30 % (w/v) PEG-600, and 10 % glycerol at 277K for FlhA_C_(R391A), reservoir solution containing 40 mM K_2_HPO_4_, 16 % (w/v) PEG-8000, and 10 % glycerol at 277K for Form 1 of FlhA_C_(R391A/V404M), and reservoir solution containing 100 mM HEPES-NaOH (pH7.5), 10 % (v/v) PEG-10,000, and 5 % 2-methyl-2,4-pentanediol at 293K for Form 2 of FlhA_C_(R391A/V404M). Each drop was prepared by mixing 1 µl of protein solution (10 mg ml^-1^) with 1 µl of reservoir solution and equilibrated to 1 ml of reservoir solution.

X-ray diffraction data were collected at the synchrotron beamline BL41XU in SPring-8 facility (Sayo-cho, Japan) with the approval of the Japan Synchrotron Radiation Research Institute (JASRI) (Proposal No. 2021A/B2739, and 2022A/B2741). The crystals were frozen in liquid nitrogen and mounted in a 100 K nitrogen stream for the X-ray diffraction data collection. The diffraction data were processed with *MOSFLM*^49^ and scaled with *AIMLESS*^50^. The initial phase was calculated by the molecular replacement method with the *Phaser-MR* in *Phenix*^51^ program using the structure of FlhAc (PDB ID: 3A5I or 6AI0) as a search model. The atomic model building and refinement were carried out using *Coot*^52^ and *Phenix*, respectively. The statistics of the diffraction data and the refinement are summarized in Supplementary Table 1. The atomic coordinates have been deposited in the Protein Data Bank with accession codes 23OT for the semi-closed from of FlhA_C_(R391A), 23PA for the semi-closed from of FlhA_C_(R391A/V404L), and 23PK for the open from of FlhA_C_(R391A/V404L).

Structural comparisons between wild-type FlhA_C_ and its mutant variants were conducted by UCSF ChimeraX^53^. All figures including structural models of FlhA_C_ were prepared using UCSF ChimeraX.

### Molecular dynamics simulations of FlhA_C_(R391A) and FlhA_C_(R391A/V404L)

For molecular dynamics (MD) simulations of FlhA_C_(R391A) and FlhA_C_(R391A/V404L), Arg-391 and Val-404 in the open-form structure of wild-type FlhA_C_ (PDB ID: 3A5I) were substituted with Ala and Leu, respectively. The AMBER ff14SB force field^54^ was applied to these mutant FlhA_C_ proteins. Each system was solvated in a cubic box of SPC/Eb water molecules^55^ containing 0.17 M KCl^56^, with a minimum distance of 12 Å between the protein and the periodic box boundaries.

All simulations were performed using the pmemd.cuda module^57^ of AMBER20/21^58^. MD simulations of FlhAC(R391A) and FlhAC(R391A/V404L) were initiated from their open conformations and independently conducted at 300 K. Following energy minimization, each system was equilibrated for 1 ns at 300 K and 1 atm with positional restraints applied to the protein main-chain atoms. Subsequently, unrestrained production MD simulations were carried out for 8 μs. The last 2 μs of the equilibrated MD trajectories were clustered into 10 groups using the K-means algorithm via ClusCo^59^. The structures representing the most populated clusters were selected for this study.

### Structural modeling

Structural models of the *Salmonella* FlhA_C_–FliK_C_ complex were generated using the AlphaFold3 prediction server^60^.

### Statistics and reproducibility

Statistical tests, sample size, and number of biological replicates are reported in the figure legends.

## Data availability

The X-ray crystal structure and structure factors of the semi-closed form of FlhA_C_(R391A) and the semi-closed and open forms of FlhA_C_(R391A/V404M) have been deposited in Protein Data Bank under accession codes 23OT, 23PA, and 23PK, respectively. All data generated during this study are included in this published article, Supplementary Information and Supplementary Data File. Strains, plasmids, polyclonal antibodies and all other data are available from the corresponding author upon reasonable request.

## Acknowledgements

We acknowledge Yasuyo Abe and Yoshie Kushima for technical assistance and beamline staffs at SPring-8 for technical help in use of beamlines BL41XU. This work was supported in part by JSPS KAKENHI Grant Numbers JP20K15749 and JP22K06162 (to M.K.), JP19H03182, JP22H02573, and JP22K19274 (to T.M.), JP16J01859 and JP23K14158 (to N.T.), JP21H01772 (to T.U.), and JP15H02386 and JP21H02443 (to K.I.). This work has also been supported by Research Support Project for Life Science and Drug Discovery (BINDS) from AMED under Grant Number JP23am121003, JP24am121003, and JP25am121003 (to K.N.), by the Cyclic Innovation for Clinical Empowerment (CiCLE) from AMED under Grant Number JP17pc0101020 (to K.N.), and by JEOL YOKOGUSHI Research Alliance Laboratories of The University of Osaka (to K.N.).

## Author Contributions

K.N. and T.M. conceived and designed research; M.K. and T.M. performed genetic, biochemical, and physiological experiments; G.S., N.T. and K.I. prepared the protein sample and performed crystallization, crystallographic data collection, processing, refinement, and model building. T.U. performed HS-AFM and analyzed the data; A.K. performed MD simulation; T.M. and K.N. wrote the paper based on discussion with other authors.

## Competing Interests

The authors declare no competing interests.

## Notes

### Competing Interest Statement

The authors have declared no competing interest.

## References

1. Minamino, T. & Kinoshita, M. Structure, assembly, and function of flagella responsible for bacterial locomotion. EcoSal Plus eesp-0011-2023 (2023).

2. Minamino, T., Doi. H. & Kutsukake K. Substrate specificity switching of the flagellum-specific export apparatus during flagellar morphogenesis in *Salmonella typhimurium*. Biosci. Biotechnol. Biochem. 63, 1301–1303 (1999).

3. Hirano, T., Minamino, T., Namba, K. & Macnab, R.M. Substrate specificity classes and the recognition signal for *Salmonella* type III flagellar export. J. Bacteriol. 185, 2485–2492 (2003).

4. Minamino, T., González-Pedrajo, B., Yamaguchi, K., Aizawa, S.-I. & Macnab, R. M. FliK, the protein responsible for flagellar hook length control in *Salmonella*, is exported during hook assembly. Mol. Microbiol. 34, 295–304 (1999).

5. Moriya, N., Minamino, T., Hughes, K. T., Macnab, R. M. & Namba, K. The type III flagellar export specificity switch is dependent on FliK ruler and a molecular clock. J. Mol. Biol. 359, 466–477 (2006).

6. Erhardt, M., Singer, H. M., Wee, D. H., Keener, J. P. & Hughes, K.T. An infrequent molecular ruler controls flagellar hook length in *Salmonella enterica*. EMBO J. 30, 2948–2961 (2011).

7. Kutsukake, K., Minamino, T. & Yokoseki, T. Isolation and characterization of FliK-independent flagellation mutants from *Salmonella typhimurium*. J. Bacteriol. 176, 7625–7629 (1994).

8. Williams, A.W., et al. Mutations in *fliK* and *flhB* affecting flagellar hook and filament assembly in *Salmonella typhimurium*. J. Bacteriol. 178, 2960–2970 (1996).

9. Fraser, G.M., et al. Substrate specificity of type III flagellar protein export in *Salmonella* is controlled by subdomain interactions in FlhB. Mol. Microbiol. 48, 1043–1057 (2003).

10. Minamino, T., Ferris, H.U., Morioya, N., Kihara, M. & Namba, K. Two parts of the T3S4 domain of the hook-length control protein FliK are essential for the substrate specificity switching of the flagellar type III export apparatus. J. Mol. Biol. 362, 1148–1158 (2006).

11. Minamino, T., Moriya, N., Hirano, T., Hughes, K.T. & Namba, K. Interaction of FliK with the bacterial flagellar hook is required for efficient export specificity switching. Mol. Microbiol. 74, 239–251 (2009).

12. Kinoshita, M., Aizawa, S.I., Inoue, Y., Namba, K. & Minamino, T. The role of intrinsically disordered C-terminal region of FliK in substrate specificity switching of the bacterial flagellar type III export apparatus. Mol. Microbiol. 105, 572–588 (2017).

13. Minamino, T., Inoue, Y., Kinoshita, M. & Namba, K. FliK-driven conformational rearrangements of FlhA and FlhB are required for export switching of the flagellar protein export apparatus. J. Bacteriol. 202, e00637–19 (2020).

14. Kinoshita, M., Tanaka, S., Y. Inoue, Y., Namba, K. Aizawa, S.I. & Minamino, T. The flexible linker of the secreted FliK ruler is required for export switching of the flagellar protein export apparatus. Sci. Rep. 10, 838 (2020).

15. Minamino, T., Kinoshita, M. & Namba, K. Insight into distinct functional roles of the flagellar ATPase complex for flagellar assembly in *Salmonella*. Front. Microbiol. 13, 864178 (2022).

16. Minamino, T., Kinoshita, M., Morimoto, Y.V. & Namba, K. Activation mechanism of the bacterial flagellar dual-fuel protein export engine. Biophys. Physicobiol. 19, e190046 (2022).

17. Minamino, T. & Namba, K. Distinct roles of the FliI ATPase and proton motive force in bacterial flagellar protein export. Nature 451, 485–488 (2008).

18. Paul, K., Erhardt, M., Hirano, T., Blair, D.F. & Hughes, K.T. Energy source of flagellar type III secretion. Nature 451, 489–492 (2008).

19. Minamino, T., Morimoto, Y.V., Hara, N. & Namba, K. An energy transduction mechanism used in bacterial type III protein export. Nat. Commun. 2, 475 (2011).

20. Minamino, T., Morimoto, Y.V., Hara, N., Aldridge, P.D. & Namba, K. The bacterial flagellar type III export gate complex is a dual fuel engine that can use both H^+^ and Na^+^ for flagellar protein export. PLoS Pathog. 12, e1005495 (2016).

21. Minamino, T., Kinoshita, M., Morimoto, Y.V. & Namba K. The FlgN chaperone activates the Na^+^-driven engine of the *Salmonella* flagellar protein export apparatus. *Commun*. Biol. 4, 335 (2021).

22. Minamino, T., Morimoto, Y.V., Kinoshita, M. & Namba, K. Membrane voltage-dependent activation mechanism of the bacterial flagellar protein export apparatus. Proc. Natl. Acad. Sci. USA 118, e2026587118 (2021).

23. Minamino, T. & Macnab, R.M. Domain structure of *Salmonella* FlhB, a flagellar export component responsible for substrate specificity switching. J. Bacteriol. 182, 4906–4919 (2000).

24. Bange, G., et al. FlhA provides the adaptor for coordinated delivery of late flagella building blocks to the type III secretion system. Proc. Natl. Acad. Sci. USA 107, 11295–11300 (2010).

25. Minamino, T., et al. Interaction of a bacterial flagellar chaperone FlgN with FlhA is required for efficient export of its cognate substrates. Mol. Microbiol. 83, 775–788 (2012).

26. Evans L.D., Poulter, S., Terentjev, E.M., Hughes, C.& Fraser, G.M. A chain mechanism for flagellum growth. Nature 504, 287–290 (2013).

27. Kinoshita, M., Hara, N., Imada, K., Namba, K. & Minamino, T. Interactions of bacterial chaperone-substrate complexes with FlhA contribute to co-ordinating assembly of the flagellar filament. Mol. Microbiol. 90, 1249–1261 (2013).

28. Xing, Q., et al. Structure of chaperone-substrate complexes docked onto the export gate in a type III secretion system. Nat. Commun. 9, 1773 (2018).

29. Qu, D., Jiang, M., Duffin, C., Hughes, K.T. & Chevance, F.F.V. Targeting early proximal-rod component substrate FlgB to FlhB for flagellar-type III secretion in *Salmonella*. PLOS Genet. 18, e1010313 (2022).

30. Bryant O.J., Dhillon, P., Hughes, C. & Fraser, G.M. Recognition of discrete export signals in early flagellar subunits during bacterial type III secretion. eLife 11, e66264 (2022).

31. Inoue, Y., Morimoto, Y. V., Namba, K. & Minamino, T. Novel insights into the mechanism of well-ordered assembly of bacterial flagellar proteins in *Salmonella*. Sci. Rep. 8, 1787 (2018).

32. Kinoshita, M., Minamino, T., Uchihashi, T. & Namba, K. FliH and FliI help FlhA bring strict order to flagellar protein export in *Salmonella*. *Commun*. Biol. 7, 366 (2024).

33. Saijo-Hamano, Y., et al. Structure of the cytoplasmic domain of FlhA and implication for flagellar type III protein export. Mol. Microbiol. 76, 260–268 (2010).

34. Abrusci, P., et al. Architecture of the major component of the type III secretion system export apparatus. Nat. Struct. Mol. Biol. 20, 99–104 (2013).

35. Kuhlen, L., Johnson, S., Cao, J., Deme, J.C. & Lea, S.M. Nonameric structures of the cytoplasmic domain of FlhA and SctV in the context of the full-length protein. PLoS One 16, e0252800 (2021).

36. Terahara, N. et al. Insight into structural remodeling of the FlhA ring responsible for bacterial flagellar type III protein export. Sci. Adv. 4, eaao7054 (2018).

37. Inoue, Y., et al. Structural insight into the substrate specificity switching mechanism of the type III protein export apparatus. Structure 27, 965–976 (2019).

38. Inoue, Y., et al. The FlhA linker mediates flagellar protein export switching during flagellar assembly. *Commun*. Biol. 4, 646 (2021).

39. Minamino, T., Kinoshita, M., Inoue, Y., Kitao, A. & Namba K. Conserved GYXLI motif of FlhA is involved in dynamic domain motions of FlhA required for flagellar protein export. Microbiol. Spectr. 10, e01110–22 (2022).

40. González-Pedrajo, B., Minamino, T., Kihara, M. & Namba, K. Interactions between C ring proteins and export apparatus components: a possible mechanism for facilitating type III protein export. Mol. Microbiol. 60, 984–998 (2006).

41. Minamino, T., et al. Roles of the extreme N-terminal region of FliH for efficient localization of the FliH-FliI complex to the bacterial flagellar type III export apparatus. Mol. Microbiol. 74, 1471–1483 (2009).

42. Hara, N., Morimoto, Y.V., Kawamoto, A., Namba, K. & Minamino, T. Interaction of the extreme N-terminal region of FliH with FlhA is required for efficient bacterial flagellar protein export. J. Bacteriol. 194, 5353–5360 (2012).

43. Notti, R.Q., Bhattacharya, S., Lilic, M., & Stebbins, C.E. A common assembly module in injectisome and flagellar type III secretionsorting platforms. Nat. Commun. 6, 7125 (2015).

44. Minamino, T., González-Pedrajo, B., Kihara, M., Namba, K. & Macnab, R.M. The ATPase FliI can interact with the type III flagellar protein export apparatus in the absence of its regulator, FliH. J. Bacteriol. 185, 3983–3988 (2003).

45. Schmiger, H. Phage P22 mutants with increased or decreased transduction abilities. Mol. Gen. Genet. 119, 75–88 (1972).

46. Ashkenazy, H., Erez, E., Martz, E., Pupko, T. & Ben-Tal, N. ConSurf 2010: calculating evolutionary conservation in sequence and structure of proteins and nucleic acids. Nucleic Acids Res. 38, W529–W533 (2010).

47. Ando, T. et al. A high-speed atomic force microscope for studying biological macromolecules. Proc. Natl. Acad. Sci. USA 98, 12468–12472 (2001).

48. Ando, T., Uchihashi, T. & Fukuma, T. High-speed atomic force microscopy for nano-visualization of dynamic biomolecular processes. Prog. Surf. Sci. 83, 337–437 (2008).

49. Battye, T.G., Kontogiannis, L., Johnson, O., Powell, H.R. & Leslie, A.G. ‘iMOSFLM: a new graphical interface for diffraction-image processing with MOSFLM’. Acta Crystallogr. D Biol. Crystallogr. 67, 271–281 (2011).

50. Evans, P.R. & Murshudov, G.N. How good are my data and what is the resoluton? Acta Crystallogr. D Biol. Crystallogr. 69, 1204–1214 (2013).

51. Adams, P.D., et al. PHENIX: a comprehensive Python-based system formacromolecular structure solution. Acta Crystallogr. D Biol. Crystallogr. 66, 213–221 (2010).

52. Emsley, P., Lohkamp, B., Scott, W.G. & Cowtan, K. Features and development of Coot. Acta Crystallogr. D Biol. Crystallogr. 66, 486–501 (2010).

53. Pettersen, E.F., et al. UCSF ChimeraX: Structure visualization for researchers, educators, and developers. Protein Sci. 30, 70–82 (2021).

54. Maier, J.A., et al. ff14SB: Improving the Accuracy of Protein Side Chain and Backbone Parameters from ff99SB. J. Chem. Theory Comput. 11, 3696–3713 (2015).

55. Takemura, K. & Kitao, A. Water model tuning for improved reproduction of rotational diffusion and NMR spectral density. J. Phys. Chem. B 116, 6279–6287 (2012).

56. Joung, I.S. & Cheatham, T.E. Determination of alkali and halide monovalent ion parameters for use in explicitly solvated biomolecular simulations. J. Phys. Chem. B 112, 9020–9041 (2008).

57. Gotz, A.W., et al. Routine microsecond molecular dynamics simulations with AMBER on GPUs. 1. generalized born. J. Chem. Theory Comput. 8, 1542–1555 (2012).

58. Case, D.A. et al. Amber 2021, University of California, San Francisco (2021).

59. Jamroz, M. & Kolinski, A. ClusCo: clustering and comparison of protein models. BMC Bioinformatics 14, 62. (2013)

60. Abramson, J. et al. Accurate structure prediction of biomolecular interactions with AlphaFold 3. Nature 630, 493–500 (2024).

