## Supplementary Information for "A single conserved residue in FlhA couples export gate activation to substrate specificity switching"

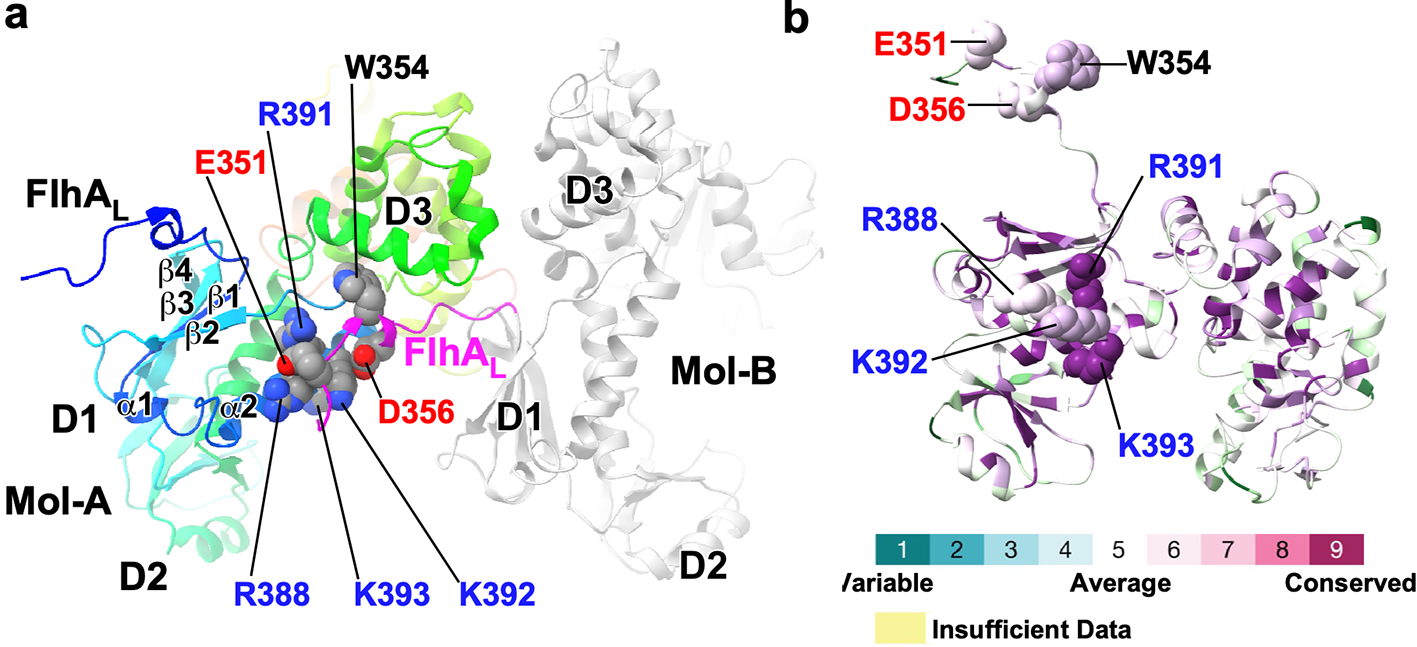

**Supplementary Fig. 1. Intermolecular FlhA_C_-FlhA_C_ interactions observed in the crystal structure. (a)** Crystal structure of the open form of the FlhA cytoplasmic domain (FlhA_C_) (PDB ID: 3A5I). The asymmetric unit contains two FlhA_C_ molecules, Mol-A (rainbow) and Mol-B (light grey). In Mol-B, two acidic residues, Glu-351 and Asp-356, located in the linker region (FlhA_L_, magenta), interact with a positively charged cluster formed by Arg-388, Arg-391, Lys-392, and Lys-393 on the surface of domain D1 of Mol-A. Trp-354 in FlhA_L_ of Mol-B binds to a hydrophobic patch at the interface between domains D1 and D3 of Mol-A. These interactions are conserved in the FlhA_C_ ring of *Vibrio parahaemolyticus* (PDB ID: 7AMY) **(b)** Evolutionary conservation of FlhA_C_ residues. Conservation scores were calculated using the ConSurf server based on FlhA sequences from 150 bacterial species and mapped onto the FlhA_C_ structure. Arg-391 and Lys-393 are highly conserved, Trp-354 and Lys-392 show relatively high conservation, whereas Glu-351, Asp-356, and Arg-388 exhibit moderate conservation.

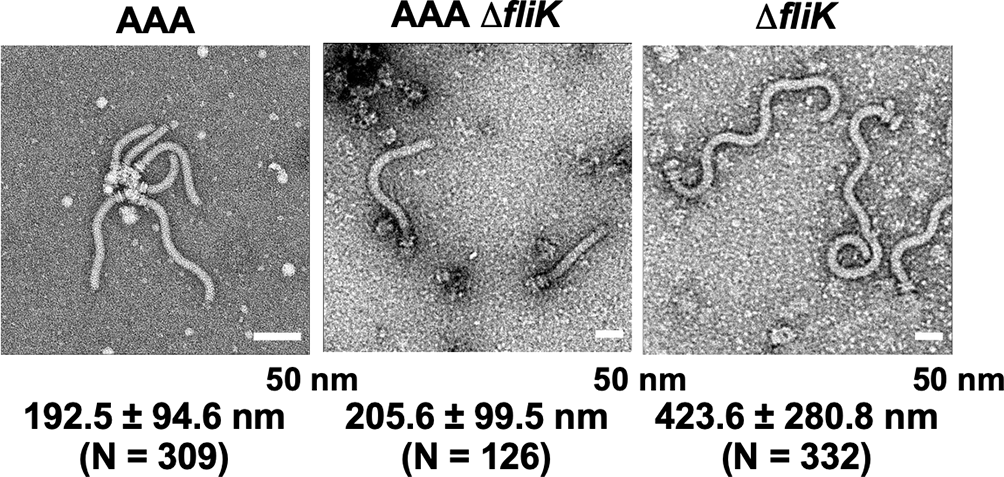

**Supplementary Fig. 2. Effect of the R391A/K392A/K393A (AAA) mutation on the hook polymerization rate.** Electron micrographs of polyhook–basal bodies isolated from *Salmonella* AAA, AAA Δ*fliK*, and Δ*fliK* mutants. The average polyhook length and standard deviation are shown. *N* indicates the number of structures measured.

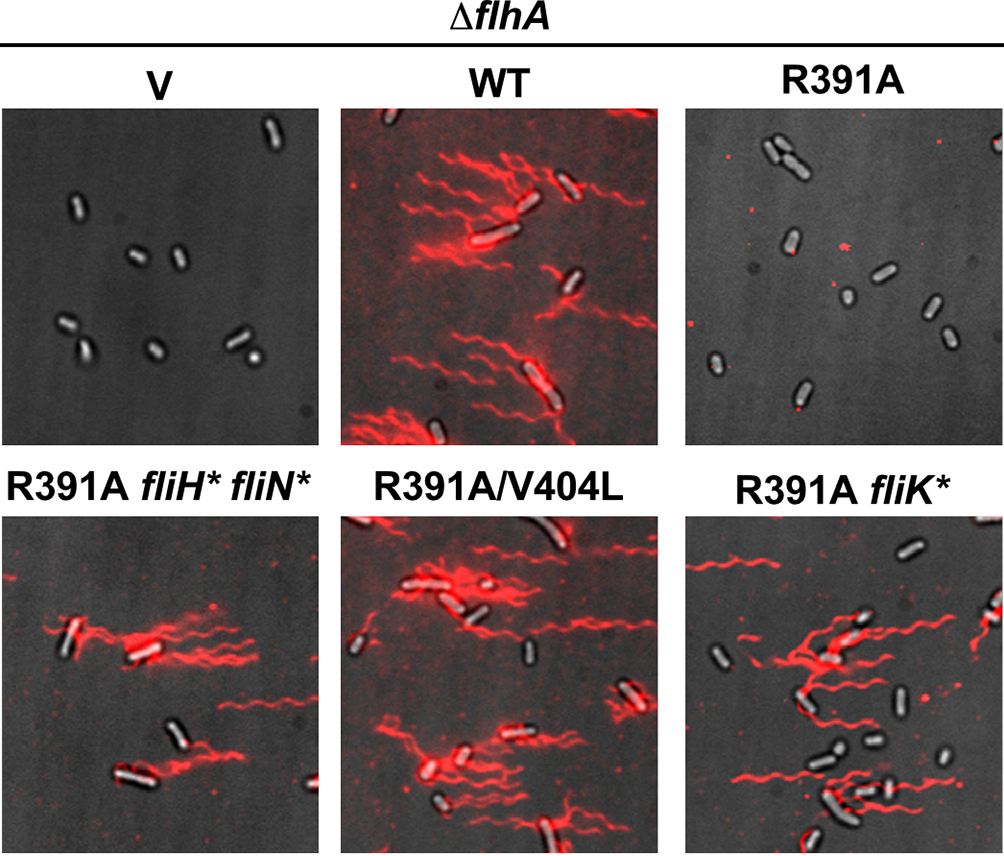

**Supplementary Fig. 3**. **Effect of the *flhA*(R391A) mutation and its suppressor mutations on flagellar filament formation.** Fluorescent images of NH001 carrying pTrc99AFF4 (V), pMM130 (WT), or pMM130(R391A), and *flhA*(R391A) suppressor strains. MMA391-6 (R391A *fliH** *fliN**) and MMA391-8 (R391A *fliK**) carry chromosomal suppressor mutations, whereas MMA391-7 carries an intragenic suppressor mutation [*flhA*(R391A/V404L)] on the plasmid. Cells were grown from fresh colonies in L-broth containing ampicillin to stationary phase, and flagellar filaments were labeled with Alexa Fluor 594. Fluorescence images of labeled filaments (red) were merged with corresponding bright-field images of the cell bodies.

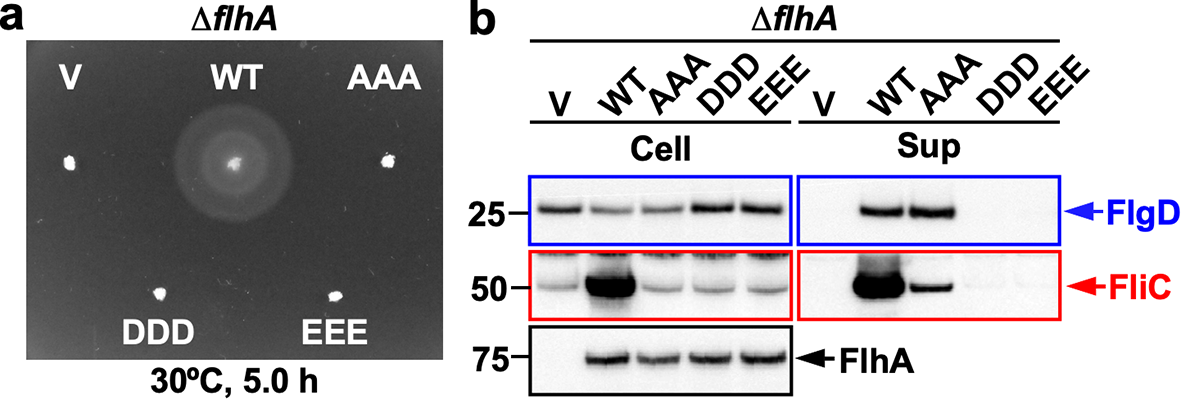

**Supplementary Fig. 4. Effect of replacement of the positively charged cluster with acidic residues.** **(a)** Soft-agar motility assays of the *Salmonella* NH001 strain (*∆flhA*) carrying pTrc99AFF4 (V), pMM130 (WT), pYI004 (AAA), pMKM130-3D (DDD), or pMKM130-3E (EEE) in soft agar. Plates were incubated at 30ºC for 5 h. At least seven independent assays were performed. **(b)** Immunoblot analysis of hook-type (FlgD) and filament-type (FliC) substrate secretion. Whole-cell proteins (Cell) and culture supernatants (Sup) were prepared from the transformants shown in (a). Hook-type and filament-type substrates are highlighted in blue and red, respectively. Molecular mass markers (kDa) are shown on the left. Three independent assays were performed.

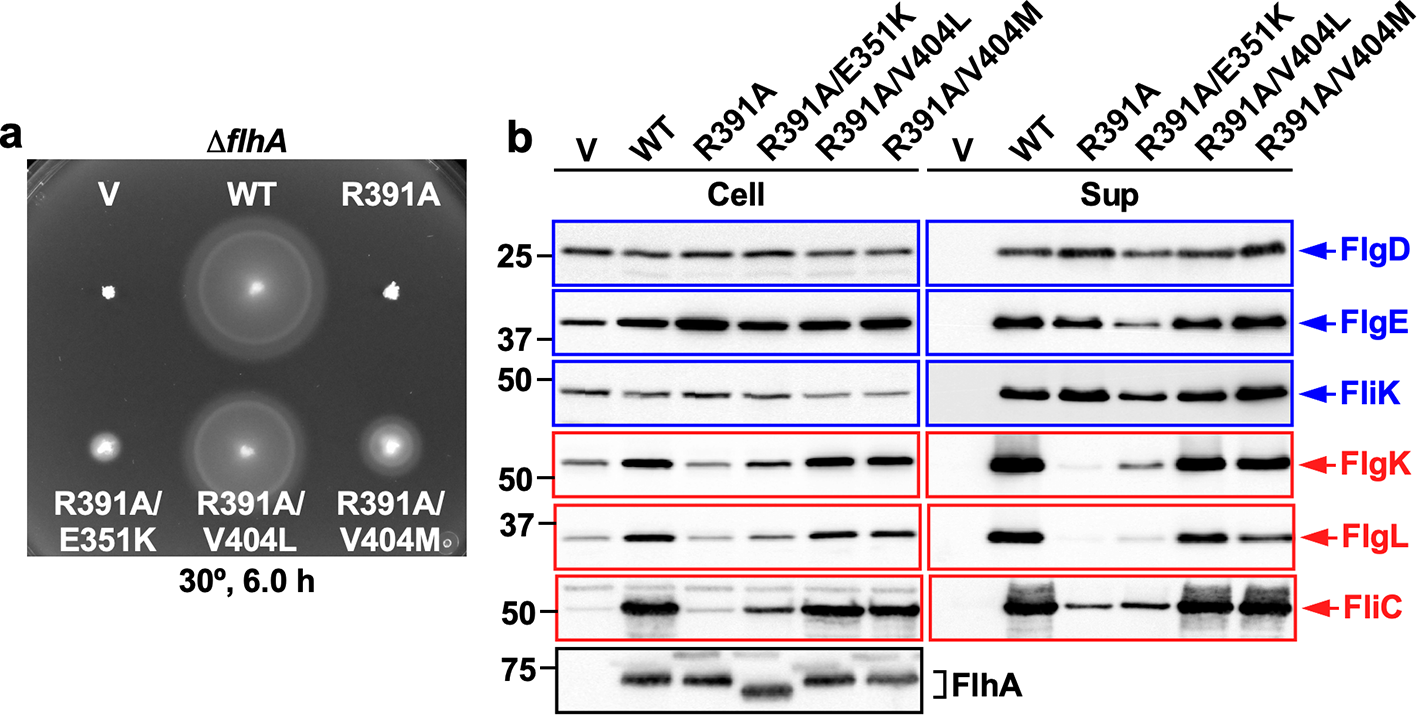

**Supplementary Fig. 5. Characterization of intragenic suppressor mutants isolated from the *flhA*(R391A) mutant. (a)** Soft-agar motility assays of the *Salmonella* NH001 strain (*∆flhA*) carrying pTrc99AFF4 (V), pMM130 (WT), pMM130(R391A), pMM130(R391A/E351K), pMKM130(R391A/V404L), or pMKM130(R391A/V404M). Plates were incubated at 30ºC for 6 h. At least seven independent assays were performed. **(b)** Secretion assays of hook-type and filament-type substrates. Whole-cell proteins (Cell) and culture supernatants (Sup) were prepared from the strains shown in (a) and analyzed by immunoblotting using the indicated polyclonal antibodies. Hook-type and filament-type substrates are highlighted in blue and red, respectively. Molecular mass markers (kDa) are shown on the left. Three independent assays were performed.

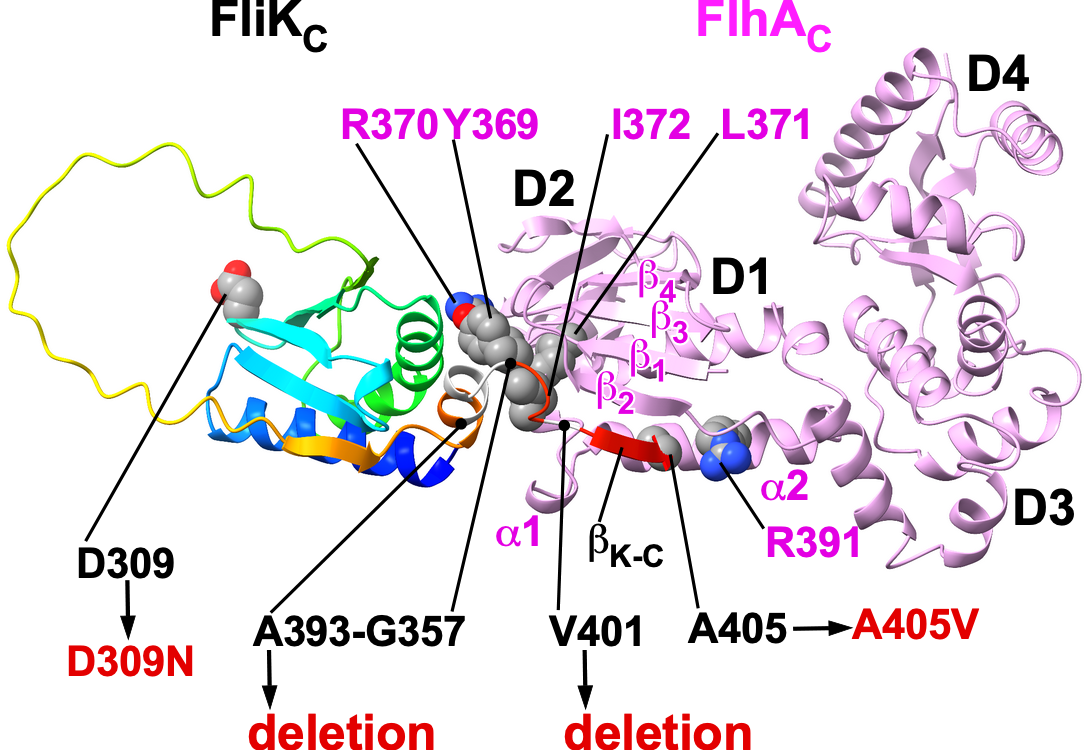

**Supplementary Fig. 6**. **Structural model of the FliK_C_–FlhA_C_ complex predicted by AlphaFold3 and mapping of extragenic suppressor mutations in FliK_C_.** 　AlphaFold3-predicted structural model of the complex between the C-terminal export-switch domain of FliK (FliK_C_, rainbow) and the C-terminal cytoplasmic domain of FlhA (FlhA_C_, orchid). Extragenic suppressor mutations in FliK_C_ (D309N, A405V, and deletions causing frameshifts that result in the addition of extra amino acids at the C terminus) are highlighted in red. Tyr-369, Arg-370, Leu-371, and Ile-372 in the conserved GYXLI motif of domain D1 of FlhA_C_, previously shown to be required for structural remodeling of the FlhA_C_ ring, are located at the interface between the core domain of FliK_C_ and FlhA_C_. The last five C-terminal residues of FliK, previously shown to be essential for substrate specificity switching of the fT3SS, adopt a β-strand (β_K-C_) that forms an antiparallel five-stranded β-sheet together with four β-strands (β1–β4) in domain D1 of FlhA_C_. Ala-405 of FliK makes hydrophobic contacts with helix α2 in domain D1 of FlhA_C_, which also contains a positively charged cluster. Notably, the *fliK*(A405V) mutation, which was isolated six times as an extragenic suppressor of the *flhA*(R391A) mutant in this study, has also been identified as an extragenic suppressor of the *flhA*(Y369A/R370A/L371A/I372A) mutant. This model suggests that both parental polyhook-causing mutations in FlhA_C_ and extragenic suppressor mutations in FliK_C_ cluster at or near the predicted FlhA_C_–FliK_C_ interface, supporting a direct role of FliK_C_ in inducing conformational remodeling of FlhA_C_ required for substrate specificity switching.

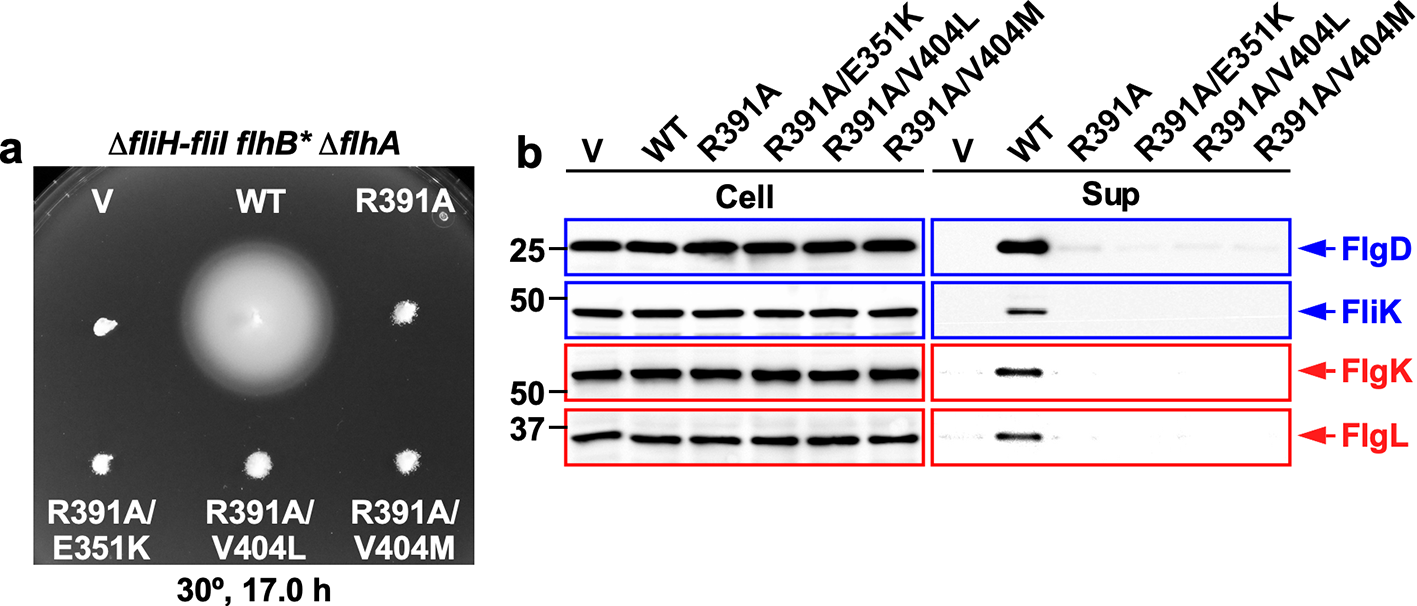

**Supplementary Fig. 7**. **Effect of FliH and FliI deletion on motility and flagellar protein export in intragenic suppressor mutants of the *flhA*(R391A) mutant. (a)** Soft-agar motility assays of the *Salmonella* NH004 (Δ*fliHI flhB** Δ*flhA*) strain carrying pTrc99AFF4 (V), pMM130 (WT), pMKM130(R391A), pMKM130(R391A/E351K), pMKM130(R391A/V404L) or pMKM130(R391A/V404M). Plates were incubated at 30°C for 17 h. At least seven independent assays were performed. **(b)** Secretion assays of hook-type and filament-type substrates in the strains shown in (a). Whole-cell proteins (Cell) and culture supernatants (Sup) were analyzed by immunoblotting using the indicated polyclonal antibodies. Hook-type and filament-type substrates are highlighted in blue and red, respectively. Molecular mass markers (kDa) are shown on the left. Three independent assays were performed.

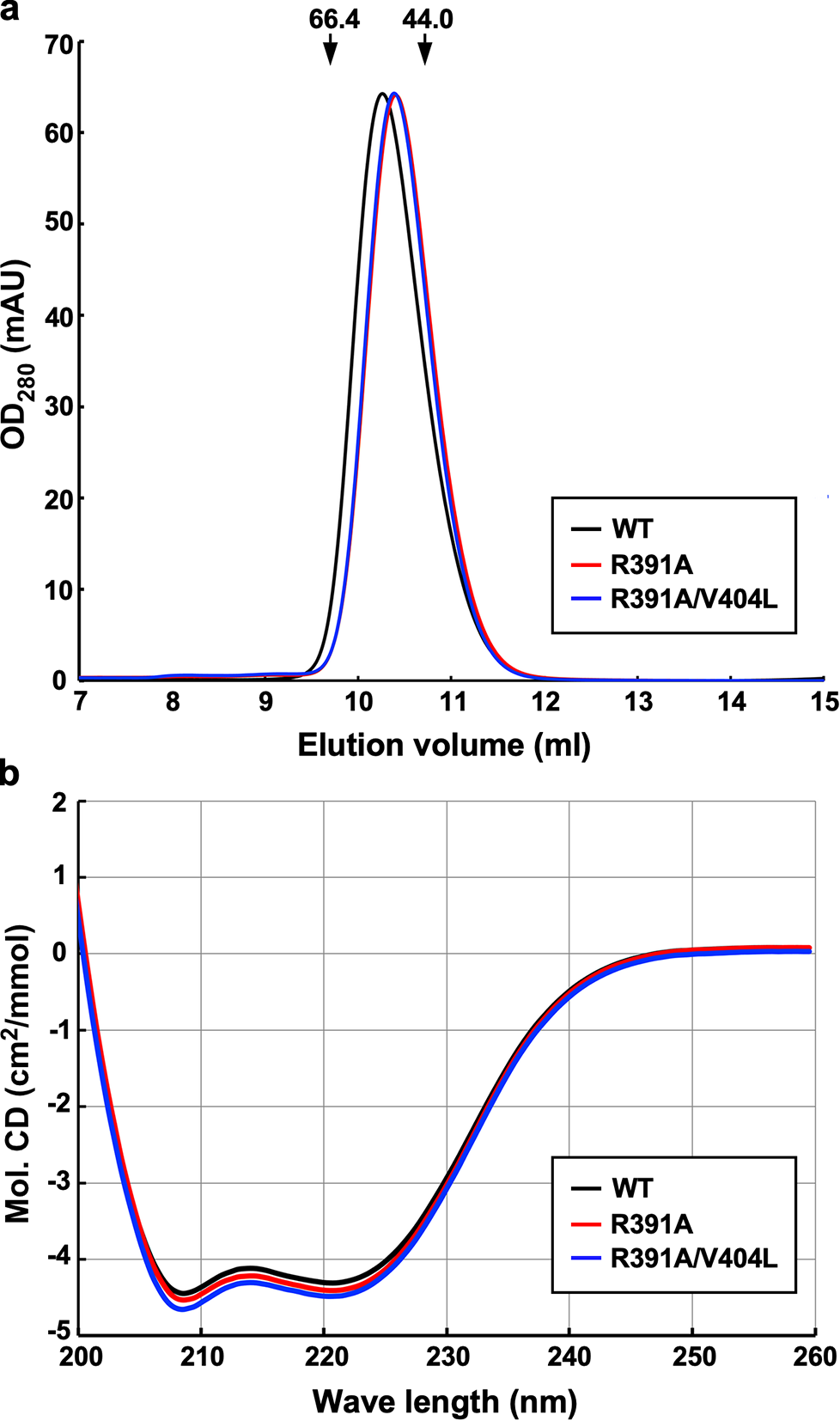

**Supplementary Fig. 8**. **Effects of the R391A and R391A/V404L mutations on the monomeric conformation of FlhA_C_. (a)** Size-exclusion chromatography analysis of purified His-FlhA_C_ (WT), His-FlhA_C_(R391A), and His-FlhA_C_(R391A/V404L). A 500 μl aliquot of each protein (10 μM) was loaded onto a Superdex 75 HR 10/300 column equilibrated with buffer containing 50 mM Tris–HCl (pH 8.0) and 150 mM NaCl. The elution peaks of His-FlhA_C_ (black), His-FlhA_C_(R391A) (red), and His-FlhA_C_(R391A/V404L) (blue) were observed at 10.3 ml, 10.4 ml, and 10.4 ml, respectively. Arrows indicate the elution positions of bovine serum albumin (66.4 kDa; 9.7 ml) and ovalbumin (44 kDa; 10.7 ml). **(b)** Far-UV circular dichroism spectra of His-FlhA_C_ (WT), His-FlhA_C_(R391A), and His-FlhA_C_(R391A/V404L). Measurements were performed at room temperature in 20 mM Tris–HCl (pH 8.0) using a quartz cuvette with a path length of 1 mm.

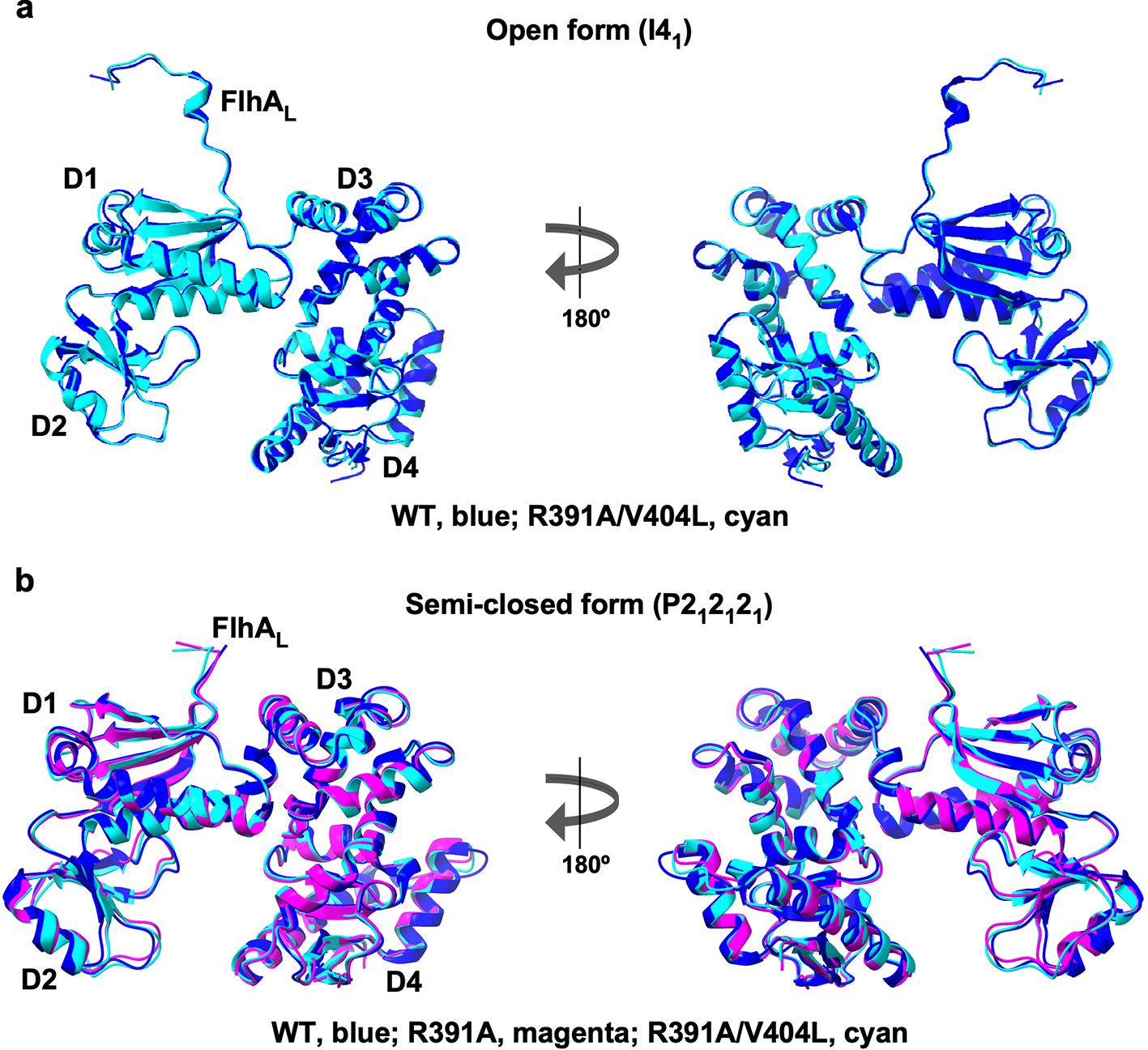

**Supplementary Fig. 9**. **Structural comparisons of FlhA_C_(R391A) and FlhA_C_(R391A/V404L) with wild-type FlhA_C_. (a)** Superposition of the open form of FlhA_C_(R391A/V404L) (PDB ID: 23PK, cyan) with that of wild-type FlhA_C_ (PDB ID: 3A5I, blue). **(b)** Superposition of the semi-closed forms of FlhA_C_(R391A) (PDB ID: 23OT, magenta) and FlhA_C_(R391A/V404L) (PDB ID: 23PA, cyan) with that of wild-type FlhA_C_ (PDB ID: 6AI0, blue).

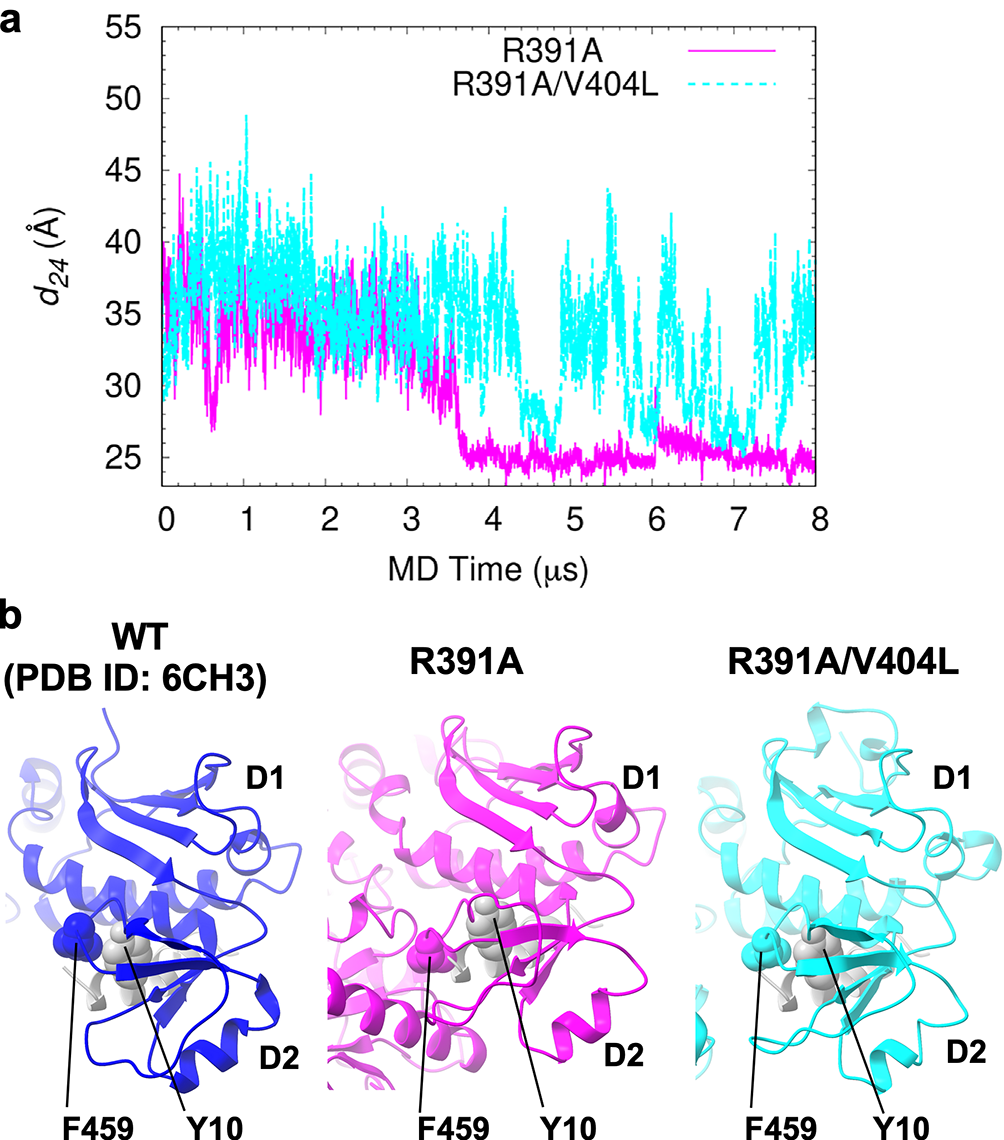

**Supplementary Fig. 10**. **Molecular dynamics (MD) simulations of FlhA_C_(R391A) and FlhA_C_(R391A/V404L). (a)** Time courses of the center-of-mass distance between domains 2 and 4 (*d_24_*) during 8-μs MD simulation. **(b)** Structural comparison of representative MD-derived structures of FlhA_C_(R391A) (magenta) and FlhA_C_(R391A/V404L) (cyan) with the crystal structure of FlhA_C_ in complex with the FliS-FliC fusion protein (PDB ID: 6CH3; FlhA_C_, blue; FliS, grey). The highly conserved Tyr-10 residue of FliS binds to a conserved hydrophobic dimple at the interface between domains D1 and D2, which includes Phe-459. Tyr-10 forms a hydrophobic contact with Phe-459 of FlhA_C_(R391A/V404L), whereas this interaction is not observed in FlhA_C_(R391A).

Supplementary Table 1. Summary of X-ray data collection and refinement statistics

|  | R391A | R391A/V404L | |
| --- | --- | --- | --- |
|  |  | Form 1 | Form 2 |
| **Data collection** |  |  |  |
| Space group | *P*2_1_2_1_2_1_ | *P*2_1_2_1_2_1_ | *I*4_1_ |
| Cell dimensions |  |  |  |
| *a*, *b*, *c* (Å) | 50.5, 95.1, 186.3 | 51.3, 92.3, 187.0 | 216.2, 216.2, 65.6 |
| α, β, γ (°) | 90.00, 90.00, 90.00 | 90.00, 90.00, 90.00 | 90.00, 90.00, 90.00 |
| Resolution (Å) | 84.7-3.20 (3.42-3.20) | 93.5-2.79 (2.94-2.79) | 68.4-3.95 (4.42-3.95) |
| *R*_merge_ | 0.132 (0.452) | 0.075 (0.475) | 0.147 (0.464) |
| *I* / *σI* | 7.4 (3.2) | 12.1 (2.6) | 7.2 (3.7) |
| CC(1/2) | 0.987 (0.893) | 0.998 (0.918) | 0.995 (0.860) |
| Completeness (%) | 97.9 (98.2) | 100 (100) | 100.0 (100.0) |
| Redundancy | 4.9 (5.2) | 4.3 (4.5) | 5.3 (5.6) |
| **Refinement** |  |  |  |
| Resolution (Å) | 84.7-3.20 (3.30-3.20) | 65.7-2.79 (2.85-2.79) | 68.4-3.95 (4.09-3.95) |
| No. reflections | 15032 | 22880 | 13583 |
| *R*_work_ / *R*_free_ | 0.236/0.264 | 0.258/0.291 | 0.215/0.251 |
| No. atoms |  |  |  |
| Protein | 5163 | 5165 | 5300 |
| Ligand/ion | - |  | - |
| Water | - |  | - |
| *B*-factors (Å^2^) |  |  |  |
| Protein | 66.8 | 67.1 | 117.0 |
| Ligand/ion | - | - | - |
| Water | - | - | - |
| Ramachandran Plot (%) |  |  |  |
| Favored | 97.1 | 95.0 | 93.3 |
| Allowed | 2.9 | 4.5 | 6.5 |
| Outliers | 0 | 0.5 | 0.3 |
| R.m.s. deviations |  |  |  |
| Bond lengths (Å) | 0.004 | 0.007 | 0.004 |
| Bond angles (°) | 0.71 | 0.97 | 0.79 |

Values in parentheses are for the highest resolution shell.

**Supplementary Table 2. Strains and plasmids used in this study**

| Strain/Plasmid | Relevant characteristics | References |
| --- | --- | --- |
| ***E. coli*** |  |  |
| BL21 Star (DE3) | Overexpression of proteins | Novagen |
| ***Salmonella*** |  |  |
| SJW1368 | ∆*cheW–flhD* | 1 |
| NH001 | ∆*flhA* | 2 |
| NH002 | *flhB(P28T)* *∆flhA* | 2 |
| NH003 | ∆*fliH-fliI ∆flhA* | 2 |
| NH004 | ∆*fliH-fliI flhB(P28T)* *∆flhA* | 2 |
| MMA391-xx | Pseudorevertants isolated from NH001 carrying pMKM130(R391A) | This study |
| **Plasmids** |  |  |
| pTrc99AFF4 | Modified pTrc expression vector | 3 |
| pGEX-6p-1 | Expression vector | GE Healthcare |
| pMKGK2 | pTrc99A/ FlgK | 4 |
| pMM104 | pET19b/ His-FlhA_C_ (residues 328–692 of FlhA) | 5 |
| pMM130 | pTrc99AFF4/ FlhA | 6 |
| pMMGN101 | pGEX-6p-1/ GST-FlgN | 7 |
| pMMJ1001 | pGEX-6p-1/ GST-FliJ | 8 |
| pYI004 | pTrc99AFF4 / FlhA(R391A/K392A/K393A) | 9 |
| pYI008 | pET15b/ His-FlhA_C_ | 9 |
| pMKM008(R391A) | pET15b/ His-FlhA_C_(R391A) | This study |
| pMKM008(R391A/V404L) | pET15b/ His-FlhA_C_(R391A) | This study |
| pMKM104(R391A) | pET19b/ His-FlhA_C_(R391A) | This study |
| pMKM104(R391A/V404L) | pET19b/ His-FlhA_C_(R391A/V404L) | This study |
| pMKM130(E351K) | pTrc99AFF4/ FlhA(E351K) | This study |
| pMKM130(R391A/E351K) | pTrc99AFF4/ FlhA(R391A/E351K) | This study |
| pMKM130(R391A) | pTrc99AFF4/ FlhA(R391A) | This study |
| pMKM130(R391A/V404L) | pTrc99AFF4/ FlhA(R391A/V404L) | This study |
| pMKM130(R391A/V404M) | pTrc99AFF4/ FlhA(R391A/V404M) | This study |
| pMKM130(K392A) | pTrc99AFF4/ FlhA(K392A) | This study |
| pMKM130(K393A) | pTrc99AFF4/ FlhA(K393A) | This study |
| pMKM130(V404L) | pTrc99AFF4/ FlhA(V404L) | This study |
| pMKM130(V404M) | pTrc99AFF4/ FlhA(V404M) | This study |
| pMKM130-3D | pTrc99AFF4 / FlhA(R391D/K392D/K393D) | This study |
| pMKM130-3E | pTrc99AFF4 / FlhA(R391E/K392E/K393E) | This study |

**Supplementary References**

1. Ohnishi, K., Ohto, Y., Aizawa, S., Macnab, R. M. & Iino, T. FlgD is a scaffolding protein needed for flagellar hook assembly in *Salmonella typhimurium*. *J. Bacteriol.* **176**, 2272–2281 (1994).
2. Hara, N., Namba, K. & Minamino, T. Genetic characterization of conserved charged residues in the bacterial flagellar type III export protein FlhA. *PLOS ONE* **6**, e22417 (2011).
3. Ohnishi, K., Fan, F., Schoenhals, G.J., Kihara, M. & Macnab, R.M. The FliO, FliP, FliQ, and FliR proteins of *Salmonella typhimurium*: putative components for flagellar assembly. *J. Bacteriol.* **179,** 6092–6099 (1997).
4. Furukawa, Y., Imada, K., Vonderviszt, F., Matsunami, H., Sano, K., Kutsukake, K. & Namba, K. Interactions between bacterial flagellar axial proteins in their monomeric state in solution. *J. Mol. Biol.* **318**, 889–900 (2002).
5. Minamino, T. & Macnab, R. M. Interactions among components of the *Salmonella* flagellar export apparatus and its substrates. *Mol. Microbiol*. **35**, 1052–1064 (2000).
6. Kihara, M., Minamino, T., Yamaguchi, S. & Macnab, R.M. Intergenic suppression between the flagellar MS ring protein FliF of *Salmonella* and FlhA, a membrane component of its export apparatus. *J. Bacteriol.* **183,** 1655–1662 (2001).
7. Minamino, T. *et al*. Interaction of a bacterial flagellar chaperone FlgN with FlhA is required for efficient export of its cognate substrates. *Mol. Microbiol.* **83**, 775–788 (2012).
8. Minamino, T. *et al*. Role of the C-terminal cytoplasmic domain of FlhA in bacterial flagellar type III protein export. *J. Bacteriol.* **192**, 1929–1936 (2010).
9. Terahara, N. *et al.* Insight into structural remodeling of the FlhA ring responsible for bacterial flagellar type III protein export. *Sci. Adv.* **4**, eaao7054 (2018).
